# Genome analysis of the Jomon dogs reveals the oldest domestic dog lineage in Eastern Eurasia

**DOI:** 10.1101/2023.09.29.560089

**Authors:** Yohey Terai, Xiayire Xiaokaiti, Jun Gojobori, Nami Arakawa, Takao Sato, Kenji Kasai, Kenichi Machida, Kyomi Yamazaki, Naomitsu Yamaji, Hitomi Hongo, Takashi Gakuhari

**Affiliations:** Research Center for Integrative Evolutionary Science, The Graduate University for Advanced Studies, Shonan Village, Hayama, Kanagawa, 240–0193, Japan; Department of Archaeology and Ethnology, Faculty of Letters, Keio University, Mita 2–15–45, Minato, Tokyo, 108–8345, Japan; Toyama Prefectural Center for Archaeological Operations, Chayamachi 206–3, Toyama, 930–0115, Japan; Toyama Cultural Foundation, Shinsogawa 4–18, Toyama, 930–0006, Japan; Faculty of Letters, Kokugakuin University, Higashi 4–10–28, Shibuya, Tokyo, 150–0011, Japan; Ichikawa City Museum of archaeology, Horinouchi 2−26−1, Ichikawa, Chiba, 272–0837, Japan; Center for the Study of Ancient Civilizations and Cultural Resources, Kanazawa University, Kakuma, Kanazawa, Ishikawa, 920–1192, Japan

**Author notes:** Corresponding authors: Hitomi, Hongo*, Takashi Gakuhari *, Yohey Terai. equally contributed (Yohey Terai, Xiayire Xiaokaiti).

## Abstract

Dog is the oldest domesticated animal that established close relationships with humans. Due to its ancient origin, when, where, and whether a single or dual domestication event occurred is still under debate. The dogs in the Jomon period (Jomon dogs) in the Japanese archipelago had little change in morphology from 10,000 to 3,000 years ago. Therefore, we expected that the ancient genome of the Jomon dogs would provide a clue to reveal the characteristics of the ancient East Asian dogs. Here, we have sequenced the genomes of three 6000-year-old Jomon dogs, one 3000-4000-year-old Jomon dog, and four late 8th century dogs excavated in Japan. Our analyses suggest that the Jomon dogs are a distinct lineage from the previously known ancient dogs and are one of the oldest among the dogs in East Eurasian lineages. In addition, the genome of the Jomon dogs contained 9.5% of the genome of Japanese wolf ancestry due to a single introgression event. We estimated the proportion of the Jomon dog lineage genome in the genomes of dogs, which indicates that the genomic composition derived from the Jomon dog lineage is one of the major sources of modern dog genomes. Furthermore, we estimated the early admixture events of dogs in East Eurasia by analyzing the ancient genomes of the Jomon dogs. Due to the admixture events, the Jomon dog-derived genome has been one of the genomic sources of a wide range of modern dogs.

## Introduction

Dog is the oldest domesticated animal that established close relationships with humans whose subsistence was still based on hunting and foraging. It is still under debate when, where, and whether a single or dual domestication event occurred, partly due to its ancient origins and rather scarce archaeological evidence (Savolainen, et al. 2002; Pang, et al. 2009; von Holdt, et al. 2010; Thalmann, et al. 2013; Wang, et al. 2013; Shannon, et al. 2015; Frantz, et al. 2016; Wang, et al. 2016; Bergström, et al. 2022).

Whole genome phylogenetic analysis of dogs and gray wolves indicates that dogs form a monophyletic group and have diverged from the internal lineage of Eurasian gray wolves (Freedman, et al. 2014; Fan, et al. 2016; Leathlobhair, et al. 2018; Gojobori, et al. 2021). A monophyletic dog group is a sister group to the Japanese wolf, indicating that the dog lineage diverged from the common ancestor of the dog and Japanese wolf (Gojobori, et al. 2021). Previous studies indicated that the dogs initially diverged into Eastern and Western Eurasian lineages (Freedman, et al. 2014; Shannon, et al. 2015; Frantz, et al. 2016; Botigué, et al. 2017; Leathlobhair, et al. 2018). Subsequently, a third lineage of dogs, the sled dog lineage, including sled dogs and the American pre-contact dogs, was reported as another major lineage (Leathlobhair, et al. 2018; Sinding, et al. 2020). The phylogenetic relationship between the sled dog lineage and the West and East Eurasian lineages has been debated (Frantz, et al. 2016; Zhang, Sun, et al. 2020). However, recent genome analysis has shown that the sled dog lineage emerged from admixture of East and West Eurasian lineages (Gojobori, et al. 2021).

The modern dingoes and New Guinea singing dogs (NGSD) are the oldest lineages in the East Eurasian lineage. The archaeological evidence showed that the arrival of dingoes in Australia was at least 3500 years ago (Milham and Thompson 1976) or 3348-3081 years ago (Balme, et al. 2018). Genome analyses suggest that the dingo and NGSD lineages have diverged from other dog lineages 8,300 (Zhang, Wang, et al. 2020) and 10,900 years ago (Bergström, et al. 2020), respectively. In the West Eurasian lineage, the modern African dogs are estimated to have diverged from European dogs 14,000 years ago (Liu, et al. 2018), with archaeological evidence dating the earliest dog in Africa at 6300-5600 BC (Mitchell 2015). These results indicate that the dingo/NGSD and African dogs, because of their isolation from the Eurasian Continent, escaped interbreeding with other Eurasian dogs and retained the genetic characteristics of the old lineages (Larson, et al. 2012; Fan, et al. 2016; Gojobori, et al. 2021).

The ancient dog genome was first determined from a Newgrange late Neolithic dog dated to 4,800 years old (Frantz, et al. 2016), followed by two individuals from Germany dating to the Early (7,000 years old) and the end of Neolithic (4,700 years old)(Botigué, et al. 2017). The analysis of these Neolithic dogs indicates that the ancient European dogs are genetically close to the West Eurasian lineage of dogs. In addition, a 9,500 years old Siberian dog is closely related to the sled dog lineage (Sinding, et al. 2020). Subsequent genome analysis of 27 ancient dog genomes indicates that at least five major lineages of dogs already existed earlier than 11,000 years ago (Bergström, et al. 2020). A study analyzing the genomes of 72 ancient wolf genomes, including individuals before and after dog domestication, showed that the entire lineage of dogs has an affinity to the wolves of Eastern Eurasia and that the genome of the Middle Eastern wolf lineage have introgressed into the dogs of the Western Eurasian lineage (Bergström, et al. 2022). In addition, East Eurasian dogs contain the genome of the ancestor of the Japanese wolf (Gojobori, et al. 2021). The most eastern Eurasian ancient dog genomes analyzed in previous studies were from the Lake Baikal area or Siberia, and no ancient dog genomes from East Asia have been reported to date.

The oldest dog excavated in the Japanese archipelago (9,300 years old) is morphologically similar to other dogs from the Jomon Period (Shigehara and Hongo 2000) (Sugihara and Serizawa 1957; Crane and Griffin 1960; Shigehara and Hongo 2000), and the oldest burial dog is 7,400-7,200 years old (Gakuhari, et al. 2015). These ages suggest that dogs were brought to the Japanese archipelago by humans about 10,000 years ago, in the Jomon period. The dogs in Jomon period (Jomon dogs) had little change in morphology throughout this period, until about 3,000 years ago (Uchiyama 2014). Therefore, Jomon dogs are presumed to be isolated and have retained the morphological characteristics of the old domestic dogs that did not interbreed with other Eurasian dogs from 10,000 to 3,000 years ago (Shigehara 1990, 1994; Komiya 1997; Shigehara and Hongo 2000; Komiya, et al. 2015). Accordingly, analysis of the ancient genome of Jomon dogs will provide a clue to reveal the characteristics of the ancient East Asian dog lineage. In this study, we extracted DNA from Jomon and late 8th century dogs in Japan and determined the genome sequences. We analyzed the ancient genome sequences with modern dogs, wolves, and Japanese wolves to show the genetic relationship among them.

## Results

We have sequenced the genomes of three 6000-year-old Jomon dogs, one 3000-4000-year-old Jomon dog (15.1-44.1 Gb: average depth of coverage, 6.2-18.2x, reference covered 11-45%), and four late 8th century Japanese dogs (15.5-98 Gb: average depth of coverage, 6.4-40.0x, reference covered 75-96%) (Table S1). Using these ancient dog genomes, we first examined the genetic relationship between the Jomon, late 8th century dogs and modern dogs, gray wolves, and Japanese wolves (Table S1).

Principal component analysis showed that each of the gray wolf and the Japanese wolf formed separate clusters from a dog cluster (Fig. 1a). The dogs formed a cluster spread along the PC2 axis, with African and Jomon dogs at each end. Late 8th century dogs and dingo/NGSD were placed in position close to the Jomon dogs (Fig. 1a). In addition, the location of European dogs was close to that of African dogs, suggesting that the PC2 axis reflected the geographic distribution of East and West Eurasia. Using the same data set, ADMIXTURE analysis divided all individuals into four genetic compositions, the gray wolf, Japanese wolf, and East and West Eurasian dogs at the lowest CV error of K=4 (Fig. 1b, Fig. S1). Jomon dogs and dingo/NGSD showed an East Eurasian dog genetic composition, while African dogs were composed of West Eurasian genetic composition (Fig. 1b).

**Figure 1.**
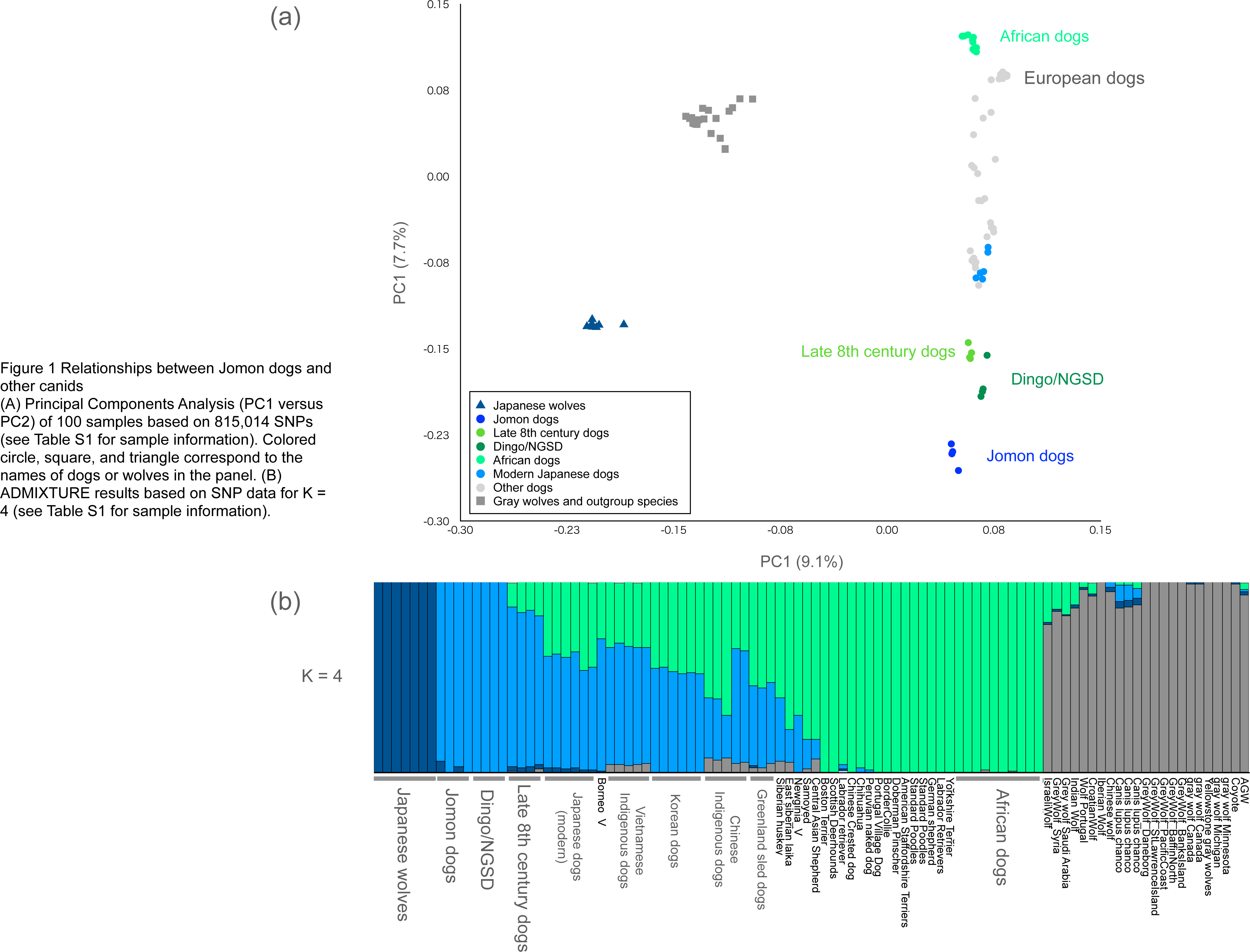
Relationships between Jomon dogs and other canids (A) Principal Components Analysis (PC1 versus PC2) of 100 samples based on 815,014 SNPs (see Table S2 for sample information). Colored circle, square, and triangle correspond to the names of dogs or wolves in the panel. (B) ADMIXTURE results based on SNP data for K = 4 (see Table S2 for sample information).

Next, we constructed phylogenetic trees based on representative dogs of three dog lineages (dingo/NGSD, African dogs, and sled dogs), plus Jomon, late 8th century dogs, gray wolves, and Japanese wolves. The dogs formed three clusters, and the Jomon and late 8th century dogs formed a monophyletic group that was a sister group to the dingo/NGSD clade (Fig. S2). Furthermore, we constructed a phylogenetic tree with ancient European and Siberian dogs. The Jomon and late 8th century dogs, together with dingo/NGSD, formed a monophyletic group of East Eurasian lineage (Fig. 2, Fig. S2). The ancient European dogs formed a monophyletic group with the dogs in West Eurasian lineage (African dogs) and the ancient Siberian dogs formed a monophyletic group with sled dogs (Fig. 2). These results indicate that Jomon dogs are members of the East Eurasian lineage and are distinct from the ancient dogs of Europe and Siberia.

**Figure 2.**
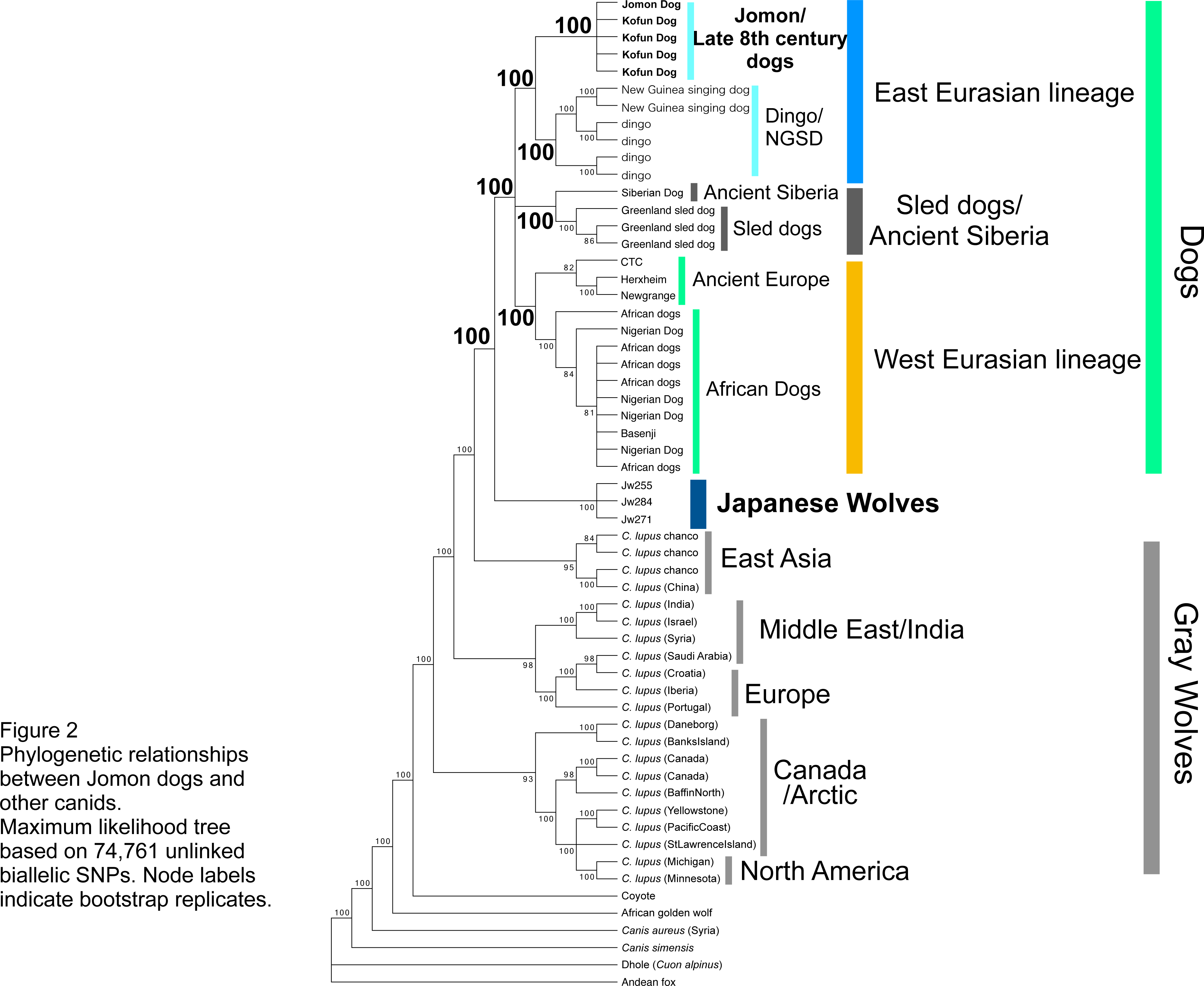
Phylogenetic relationships between JJomon dogs and other canids. Maximum likelihood tree based on 74,761 unlinked biallelic SNPs. Node labels indicate bootstrap replicates.

In Eurasia, the genomes of dogs of Eastern and Western Eurasian lineages have been admixed (Gojobori, et al. 2021). However, dingo/NGSD and African dogs rarely admixed with Eurasian dogs, and they were reported as genetically the most distantly related pair of dogs (Gojobori, et al. 2021). Therefore, if Jomon dogs belong to one of the oldest lineages of dogs in East Eurasian lineage, we expect that Jomon dogs are genetically the most distantly related to African dogs. We then examined the genetic relationship between Jomon dogs and other dogs. The *f3* statistic for all dog combinations showed that a Jomon dogs and African dogs pair have the lowest genetic affinity (Fig. S3), suggesting that Jomon dogs belong to one of the oldest lineages in the East Eurasian dogs and are the least admixed with the West Eurasian dogs. Next, we examined dogs with high genetic affinity to Jomon dogs using *f3* statistics. Late 8th century dogs had the highest genetic affinity to Jomon dogs, followed by dingo/NGSD and modern Japanese dogs (Fig. S4-S6), indicating that late 8th century dogs are the direct descendants of the Jomon dogs. No ancient dog had a higher affinity with the Jomon dogs than late 8th century dogs and dingo/NGSD (Fig. S5 and S6). These results indicate that the Jomon dogs are a distinct lineage from the previously known ancient dogs and are one of the oldest among the East Eurasian dog lineages.

A recent study reported that the genomes of East Eurasian dogs contain the genome of the Japanese wolf ancestry resulted in high genetic affinity of these two groups (Gojobori, et al. 2021). To further examine the relationship of the Jomon dogs to wolves, we analyzed the genetic affinity between Jomon dogs and gray wolves using *f3* statistics. As the result, the Japanese wolf showed the highest affinity to Jomon dogs among modern wolves (Fig. S7). We added ancient wolves to the analysis, and showed that no ancient wolf showed a higher affinity to Jomon dogs than the Japanese wolf, either (Fig. 3a). We next examined the gene flow between Jomon dogs and gray wolves including the Japanese wolf using the *f4* statistic. We found that the Japanese wolf had the highest *f4* value (Fig. S8), indicating a gene flow between Jomon dogs and the Japanese wolf. Furthermore, the *f4*-ratio revealed that the genomes of Jomon dogs contained the highest proportion of the Japanese wolf ancestry genome (9-11%), followed by late 8th century dogs and NGSD (5.0-6.7%) (Fig. 3b). The dog genome in the Japanese wolf genome was under the detection limit (Fig. S9). These results suggest a genomic introgression event from the Japanese wolf to East Eurasian dogs.

**Figure 3.**
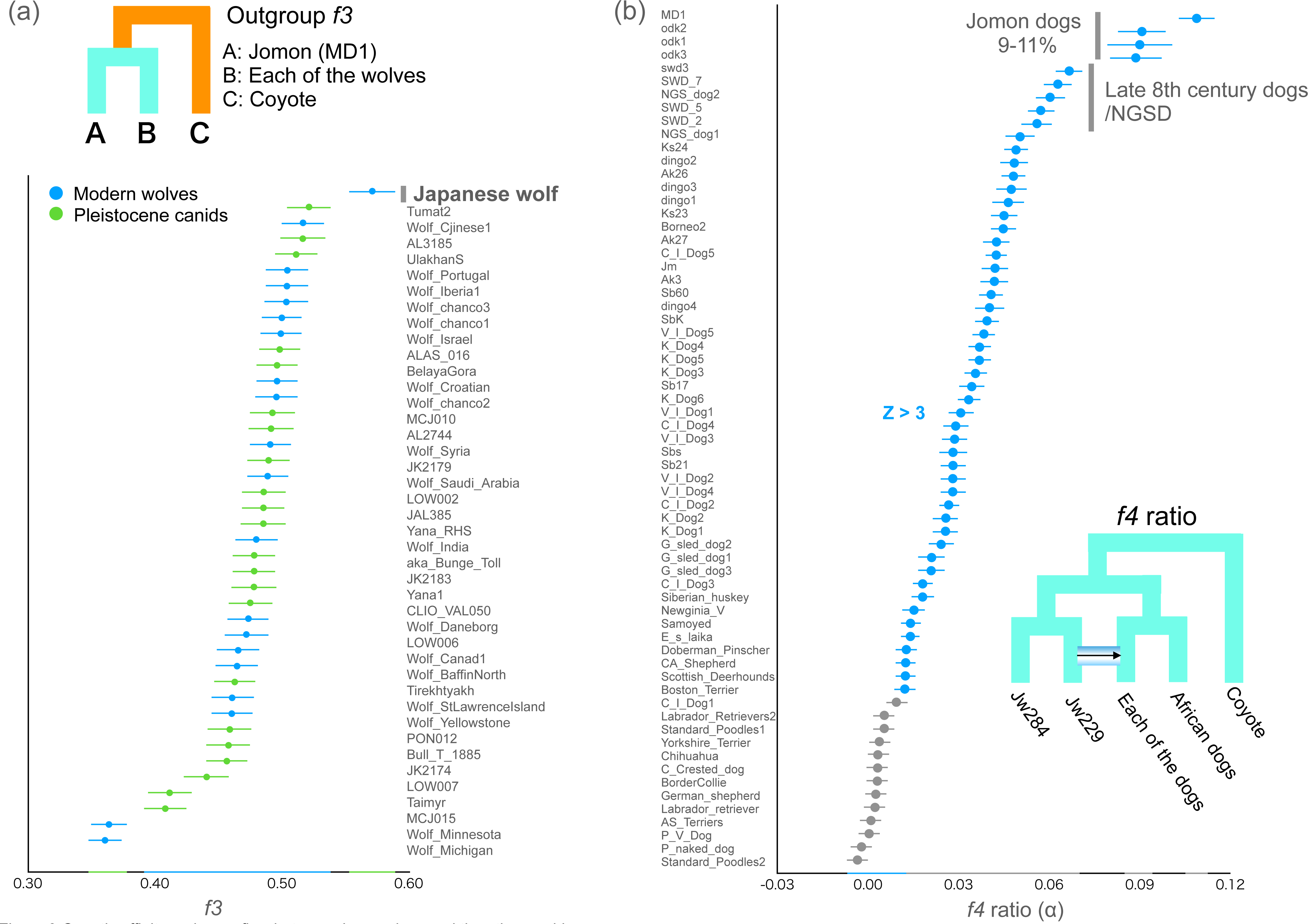
Genetic affinity and gene flow between Jomon dogs and the other canids (a) Shared genetic drift between Jomon dogs and gray wolves measured by outgroup *f3* statistics. Each of all wolves and a Jomon dog (MD1) were used as populations. Each *f3* statistical value is plotted in order of highest to lowest value from the top, and the names of the wolves are shown on the right side of the panel. Error bars represent standard errors. (b) *f4*-ratio test to estimate proportion of genome introgression from the Japanese wolf to dogs. Each f4-ratio α value is plotted in order of highest to lowest value from the top, and the names of the dogs are shown on the left side of the panel. Error bars represent standard errors. Z score above 3 is colored in blue.

We further examined the number of introgression events (waves) from wolves to dogs by qpwave. Setting the Japanese wolf, North American and Eurasian gray wolves as genomic sources, qpwave analysis indicates that two or more introgression events have occurred from wolves to dogs (Table 1: Test 1). When we excluded the Japanese wolf from the genomic sources of wolves, the introgression events were estimated to have occurred once or more (one reduction) (Test 2). These results suggest one introgression event from the Japanese wolf to a dog. When we set only East Eurasian dogs as the dog population, the analysis showed one or more introgression event(s) from wolves to East Eurasian dogs (Test 3). The introgression event was eliminated when we excluded the Japanese wolf (Test 4) or a Jomon dog (Test 5) from this analysis. This elimination of the event indicates an introgression event from the Japanese wolf (ancestry) to the Jomon dogs (ancestry). Similarly, we analyzed the introgression event from wolves to West Eurasian lineage (African dogs) (Test 7, 8) and showed an introgression event from an Israeli wolf (ancestry) to African dogs (ancestry) as reported in (Bergström, et al. 2022).

**Table 1.** The results of qpwave.

|  | Left (dogs) populations | Right (wolves) populations | p < 0.05 | Number of admixture events |
| --- | --- | --- | --- | --- |
| **Test 1** (Eastern and Western dogs) | Jomon, NGSD1, Dingo1, African1, Nigerian1 | Jw284, chanco1, Chinese1, Portugal, Israel, Michigan | rank = 0, 1 | ≥ 2 |
| **Test 2** (Eastern and Western dogs) | Jomon, NGSD1, Dingo1, African1, Nigerian1 | chanco1, Chinese1, Portugal, Israel, Michigan | rank = 0 | ≥ 1 |
| **Test 3** (Eastern dogs) | Jomon, NGSD1, NGSD2, Dingo2, Dingo4 | Jw284, chanco1, Chinese1, Portugal, Israel, Michigan | rank = 0 | ≥ 1 |
| **Test 4** (Eastern dogs) | Jomon, NGSD1, NGSD2, Dingo2, Dingo4 | chanco1, Chinese1, Portugal, Israel, Michigan | - | - |
| **Test 5** (Eastern dogs) | NGSD1, NGSD2, Dingo2, Dingo4 | Jw284, chanco1, Chinese1, Portugal, Israel, Michigan | - | - |
| **Test 6** (Eastern dogs) | Jomon, NGSD1, NGSD2, Dingo2, Dingo4 | Jw284, chanco1, Chinese1, Portugal, Michigan | rank = 0 | ≥ 1 |
| **Test 7** (Western dogs) | African1, African2, Nigerian1, Nigerian2 | Jw284, chanco1, Chinese1, Portugal, Israel, Michigan | rank = 0 | ≥ 1 |
| **Test 8** (Western dogs) | African1, African2, Nigerian1, Nigerian2 | Jw284, chanco1, Chinese1, Portugal, Michigan | - | - |
| **Test 9** (Western dogs) | African1, African2, Nigerian1, Nigerian2 | chanco1, Chinese1, Portugal, Israel, Michigan | rank = 0 | ≥ 1 |

If genomic introgression event from the Japanese wolf ancestry to the Jomon dog lineage occurred once, we can assume that the ratio of the genome of Jomon dog ancestry to that of the Japanese wolf ancestry was constantly maintained in the genomes of the other dogs admixed with Jomon dog lineage (Fig. S10). To test this possibility, the genetic affinities of Jomon dogs to each of the other dogs and that of Japanese wolves to each of the other dogs were plotted on the x-axis and y-axis, respectively. The plots showed a positive correlation between the genetic affinity to Jomon dogs and that of Japanese wolves in all dogs (Fig. 4). Thus, each dog has a constant ratio of Jomon dog ancestry to the Japanese wolf ancestry, suggesting a single introgression event from the Japanese wolf ancestry to the ancestor of the Jomon dogs.

**Figure 4.**
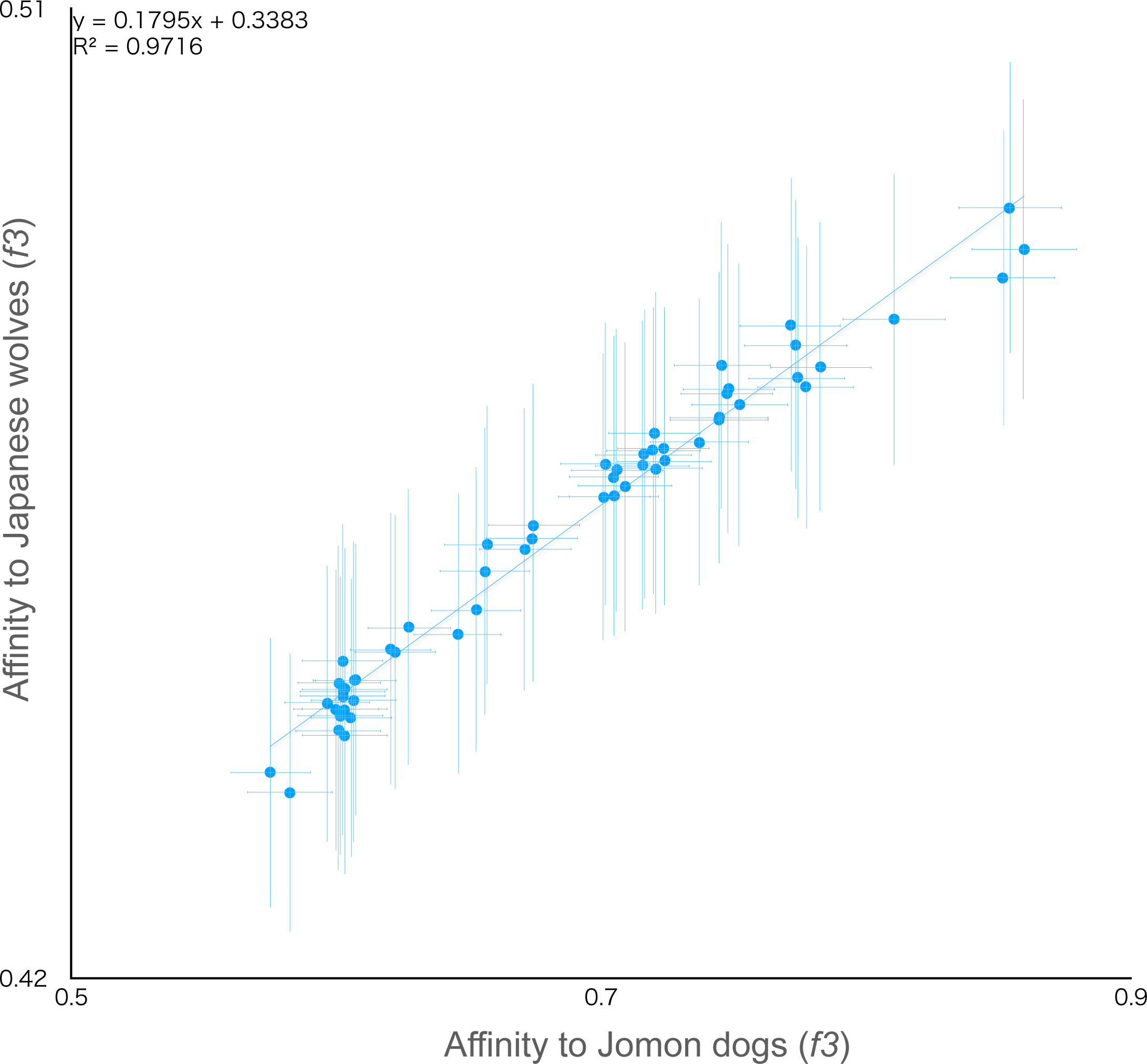
Positive correlation between genetic affinity of Japanese wolf and Jomon dog to other dogs *f3* statistics testing whether dogs share more alleles with Jomon dogs (x-axis) or Japanese wolf (y-axis). Dots show the *f3* statistics, and horizontal and vertical error bars represent standard errors. Each of the Japanese wolves and Jomon dogs individuals were used as populations.

Next, we examined the genetic relationship between Jomon dogs and dingo/NGSD, an old lineage in East Eurasia. The proportion of the West Eurasian dogs (European dogs) genome in each dog showed that most dogs, including the late 8th century dogs, contained genomes from the West Eurasian dogs (Fig. S11). However, the two dingo/NGSD individuals were below the detection limit (Z score < 3) (Fig. S11). Thus, similar to Jomon dogs, which are the most distantly related to African dogs, the genomes of dingo/NGSD contain no or little proportion of the genome of the West Eurasian dogs. Nevertheless, the proportion of the Japanese wolf ancestry in the genome differs between Jomon dogs at about 9.5% (average of four Jomon dogs) and dingo/NGSD at about 5%. This result suggests a possibility that there was an East Eurasian lineage without including the Japanese wolf ancestry genome (No-Jw-dogs) and that dingo/NGSD may have resulted from an admixture of the Jomon dog lineage and the No-Jw-dogs lineage.

To test this possibility, we plotted the genetic affinities of dingo/NGSD to each dog on the X-axis and those of Jomon dogs (Fig. S12a) or the Japanese wolf (Fig. S12b) to each dog on the Y-axis. Both graphs show Y-shaped plots, indicating that the East Eurasian dogs were separated into two groups, individuals with higher affinity to dingo/NGSD and individuals with higher affinity to Jomon dogs (Fig. S12a). Furthermore, individuals with a high affinity to Jomon dogs also have a high affinity to the Japanese wolf (Fig. S12b). This result was consistent with the positive correlation between the ratio of the Jomon dog and the Japanese wolf ancestry compositions in the genome (Fig. 4). Therefore, individuals with a higher affinity to dingo/NGSD are estimated to contain more genomes derived from No-Jw-dogs. Using qpgraph, we compared the likelihood scores between a simple dingo/NGSD and Jomon dog lineages divergence model (Likelihood score: 80.2) (Fig. S13d) and a dingo/NGSD admixture model (Likelihood score: 29.0) (Fig. S13e). The latter model was supported by a lower Likelihood score and suggests an existence of No-Jw-dogs in East Eurasia.

The sled dog lineage may have arisen from an admixture of Eastern and Western Eurasian lineages (Gojobori, et al. 2021). However, the East Eurasian source of the admixture is still unknown. To confirm the admixture event and identify this East Eurasian source, we compared the likelihood scores by qpgraph between models in which sled dog lineage has diverged a) simultaneously with the divergence of Eastern and Western Eurasian lineages, b) from the Western Eurasian lineage, c) from the dingo/NGSD lineage, d) from the Jomon dog lineage, or e-g) have arisen by the admixture of the Eastern and Western Eurasian lineage (Fig. S14). The results showed that models from an admixture of Eastern and Western Eurasian lineages had lower scores than the others. Among the models of the Eastern and Western Eurasian lineages admixture (Fig. S14e-g), the dingo/NGSD and Western Eurasian lineages admixture model (Fig. S14g) was the lowest likelihood score, suggesting that the sled dog lineage was likely to be an admixture of these two lineages.

## Discussion

### Jomon dog lineage is one of the genomic sources of the modern dogs

Dogs form a monophyletic group and are, therefore, genetically monophyletic origin (Freedman, et al. 2014; Fan, et al. 2016; Leathlobhair, et al. 2018; Gojobori, et al. 2021). This does not indicate a single origin of dog domestication, because the domestication process would have been initiated with the association of dogs with humans (Larson, et al. 2012). A single genetic origin of dogs suggests that the majority of the dog genomes would be derived from the common ancestor of the monophyletic dog group. The genomes supplied by introgressions are also the source of the modern dog genome, because the West and East Eurasian lineage of dogs contain the genome of Middle Eastern wolves and the Japanese wolf ancestry genome, respectively (Gojobori, et al. 2021; Bergström, et al. 2022).

In this study, we estimated a single event of genomic introgression of about 9.5% from the Japanese wolf ancestry into the Jomon dog lineage. Therefore, the proportion of the Japanese wolf ancestry genome is a good indicator to estimate how much of the Jomon dog lineage genome is included in the genomes of dogs. In other words, the genomic content of the Jomon dog lineage in a particular dog genome can be estimated from the proportion of the Japanese wolf ancestry genome. For example, the NGSD individuals contain 5.0-6.0% Japanese wolf ancestry genome, and these values allow us to estimate that 53-63% of the NGSD genomes are derived from the genome of the Jomon dog lineage. Using this model, we estimated the proportion of the Jomon dog-derived genome and found that modern dogs from a broad geographic region contained Jomon dog-derived genome; 45-52% in Akita and Kishu dogs, 22-25% in Greenland sled dogs, and 13% in several European dogs (Fig. 5). These proportions indicate that the genome derived from the Jomon dog lineage is one of the major sources of the modern dog genomes.

**Figure 5.**
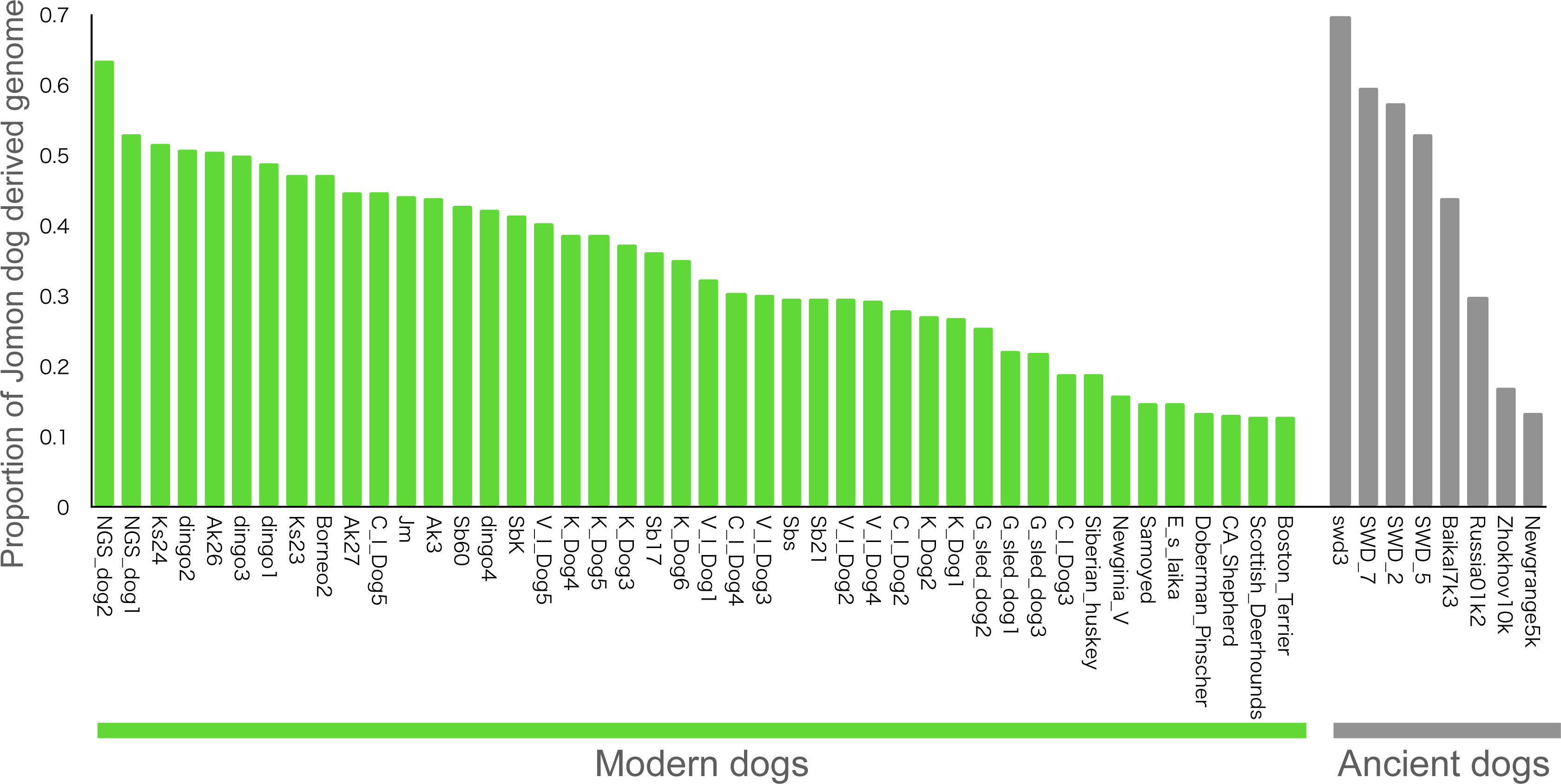
Estimated proportion of Jomon dog derived genome in dogs.

In ancient dogs, we estimated the proportion of the Jomon dog-derived genome as 53-70% for late 8th century dogs, 44% for Baikal 7k, 17% for Zhokhov 10k, and 14% for Newgrange 5k, indicating that even ancient European dogs already contained the Jomon dog genome. Since the 9,500 years old Zhokhov 10k also contains the Jomon dog-derived genome, the Jomon dog lineage was already admixed with other dogs before 9,500 years ago.

### Admixture events of dogs in East Eurasia in the Pleistocene

Previous studies of ancient dog genomes have shown that five major dog lineages existed 11,000 years ago (Bergström, et al. 2020). However, ancient genomic studies of dogs used individuals from West Eurasia and Siberia and did not use East Eurasian dogs (Frantz, et al. 2016; Botigué, et al. 2017; Bergström, et al. 2020). Therefore, the ancient admixture event of dogs in East Eurasia, the possible origin of the dog lineage, remains to be explored.

Based on the analyses in this study, we estimated the following events during the early stages of the dog evolution: 1) divergence of the dog lineage from the common ancestor of the Japanese wolf and dog, 2) divergence of the East and West Eurasian lineages, 3) divergence of the East Eurasian lineage into two lineages, 4) introgression of the Japanese wolf ancestry genome into the Jomon dog lineage, 5) admixture of the two East Eurasian lineages, arisen dingo/NGSD lineage, 6) admixture of the dingo/NGSD lineage and the West Eurasian lineage, arisen the sled dog lineage (Fig. 6). The first and fourth events were proposed to have occurred in East Eurasia (not in the Japanese archipelago)(Gojobori, et al. 2021). The divergence of the East Eurasian lineage (Event 3) and the admixture of the two East Eurasian lineages (Event 5) are both related to East Eurasian and, therefore, likely occurred in East Eurasia. Hence, the early events 1-5 in the evolution of the dog most likely occurred in East Eurasia. The sled dog lineage that appeared at the end of our estimation includes the 9,500 years old Zhokhov 10k, suggesting that the dingo/NGSD lineage and the West Eurasian lineage had admixed earlier than 9,500 years ago. Thus, the evolutionary events estimated here must have occurred during the final Pleistocene or the beginning of Holocene.

**Figure 6.**
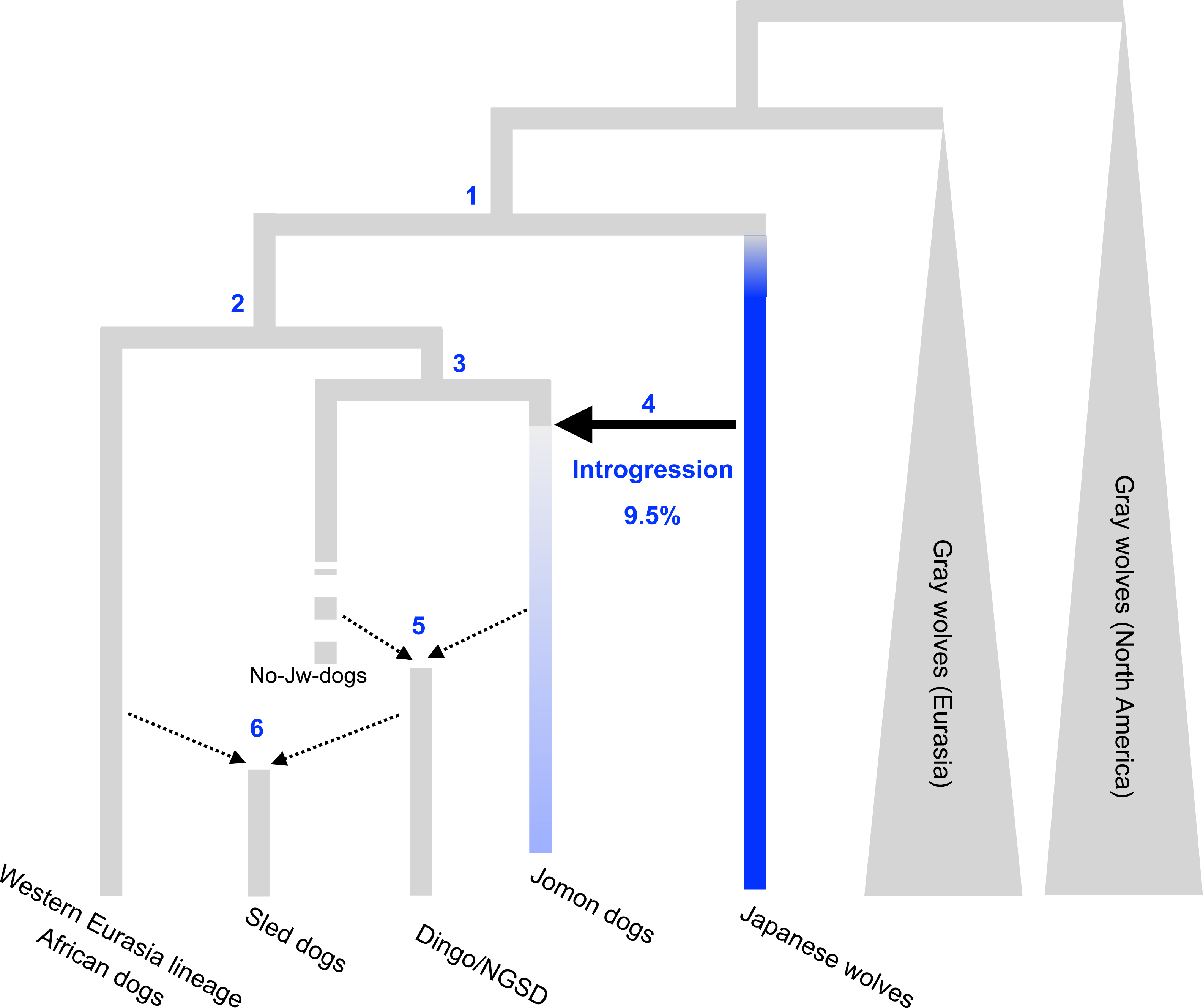
A model of the early stages of the dog evolution. Each event was supported by following resuts; event 1), phylogenomic analyses (Fig.2, S2): 2) phylogenomic analyses (Fig.2, S2), 3) *f4* ratio test (Fig. 3b, S11), *f3* biplots (Fig. S12), and qpgraph (Fig. S13): 4) *f4* ratio test (Fig. 3b, S9) and qpgraph (Fig. S13), 5) *f4* ratio test (Fig. 3b, S11) and qpgraph (Fig. S13): 6) *f4* ratio test (Fig. 3b) and qpgraph (Fig. S14). Arrow indicate introgression or admixture events.

It is unknown if humans were involved in these admixture events of dogs. However, when the dogs appeared in the Japanese Archipelago in Jomon period, they were already distinct from the wolves both in their morphology and size, suggesting that they had already been domesticated. Therefore, the ancestors of the Jomon dog lineage were likely to have been domesticated when they were still in the Eurasian Continent.

The gene flow from the Southeast Asian dog ancestry to the ancestor of the ancient European dogs before 7,000 years ago (Botigué, et al. 2017) suggests ancient East and West Eurasian dog lineages interactions. Indeed, the late 8th century dogs already contained 15-22% of the West Eurasian lineage genome, and therefore the dog that arrived in Japan in the period after the end of the Jomon period (3,000 years ago) may have been a mixture of the East and West Eurasian dog lineages.

In this study, we estimated the early admixture events of dogs in East Eurasia by analyzing the ancient genomes of the Jomon dogs. Due to the admixture events, the Jomon dog-derived genome have been one of the genomic sources of a wide range of modern dogs. Since the Jomon dog lineage is an old lineage isolated in the Japanese archipelago from 10,000 to 3,000 years ago, the Jomon dogs are the best target for estimating initial evolutionary events of dogs in East Eurasia. Thus, in the future, more detailed analyses of the early divergence of dog lineages in East Eurasia through ancient genome analysis will require analysis of bones earlier than 10,000 years old.

## Materials and Methods

### Samples, DNA extraction, and sequencing

For the Jomon dogs (odk1, odk2, odk3, and MD1) and late 8th century dogs (SWD2, SWD3, SWD5, and SWD7), the petrous bones were cut with diamond cutting disc. After washed by ultrapure water and 99.9% Ethanol, the cut-out pieces were irradiated UV-light for 30 min. Approximately 50∼100 mg bone powder was obtained from the inner part of petrous bones by drilling. DNA was extracted followed by a modified protocol of Gamba et al. 2014 as described in Gakuhari et al. 2020 (Gamba, et al. 2014; Gakuhari, et al. 2020). The powder was first digested with pre-digestion buffer containing 0.5 M EDTA, 0.65 U/ml recombinant Proteinase K, and 10% N-Laurylsarcosyl and incubate at 50°C for 15 min with shaking (900 rpm). The reaction solution was centrifuged at 13,000 g for 10 min, and the supernatant was stored (Pre-digestion supernatant). The lysis buffer containing 20 mM Tris HCl(pH 7.5), 0.7% N-lauroylsarcosine, 47.5 mM EDTA (pH 8), 0.65 U/ml recombinant Proteinase K was added to the pellet, and then the reaction solution was incubated 16 hours at 37°C with shaking (900 rpm). The reaction solution was centrifuged at 13,000 g for 10 min, and the supernatant was diluted with 3 ml TE (pH 8.0) and centrifuged using a filtration tube (Amicon® Ultra-4 Centrifugal Filter Unit 30K) at 2,000 g until final volume of 100 µl.

The DNA solution was purified by MinElute PCR Purification Kit (QIAGEN) following the instruction with pre-heated (60°C) EB buffer containing 0.05% Tween 20. Extraction of ancient DNA was performed in an ancient DNA clean room at Kanazawa university. Before sampling, all the bones were irradiated UV-light for 30 min to remove surface contaminants. We targeted the inner part of petrous bones which is recognized as optimal substrates for ancient studies. (Pinhasi et al. 2015) Before library construction, the DNAs of one Jomon dog (MD1) and three late 8th century dogs (SWD_2, SWD_5, and SWD_7) were treated by USER enzyme (NEB) for uracil removal. DNA libraries were constructed from 1 ng of genomic DNA with NEBNext Ultra II DNA Library Prep Kit for Illumina and NEBNext Multiplex Oligos for Illumina (New England Bio Labs, MA, USA). Paired-end (2 × 150 bp) sequencing was performed on the Illumina HiSeq X or NovaSeq 6000 platforms.

### Extraction of SNPs and vcf file preparation

We downloaded sequencing data of modern dogs, modern gray wolves, ancient dogs, ancient canids, and outgroup species from the database (Table S2). Sequence reads from the genomic DNA libraries of Jomoin and the late 8th century dogs (Table S1) as well as samples from the database (Table S2) were trimmed to remove nucleotides with base qualities lower than 35 on average of a 150 bp read (sum of the base-calling error probability < 0.05 in a 150 bp read) and adaptor sequences using CLC Genomics Workbench (https://www.qiagenbioinformatics.com/). The trimmed reads were mapped to the dog reference genome (CanFam3.1) using CLC Genomics Workbench. Reads showing high similarity (> 90% in > 90% of read length) were mapped to the reference genome sequences to avoid mapping the low similarity reads. Reads mapped to more than one position were removed (“ignore” option for reads mapped to multiple positions) to prevent mapping to non-unique regions. The mapping data was exported in bam file format and sorted and indexed using samtools (Li, et al. 2009). The duplicated reads in bam files were marked by the MarkDuplicates algorithm implemented in GATK v4.2 (https://gatk.broadinstitute.org/hc/en-us). We performed genotype calling on all individuals analyzed in this study using the HaplotypeCaller algorithm in GATK v4.2. Genotypes of all individuals were output as gvcf format (-ERC GVCF option). All gvcf files were combined into a single gvcf format file by the CombineGVCFs algorithm in GATK v4.2. The combined file was genotyped by the GenotypeGVCFs algorithm and filtered by Filtervcf in GATK v4.2 with parameters; --filter-expression “QD < 2.0” --filter-name “QD2” --filter-expression “QUAL < 30.0” --filter-name “QUAL30” --filter-expression “FS > 200.0” --filter-name “FS200” --filter-expression “SOR > 10.0” -filter-name “SOR10” --filter-expression “ReadPosRankSum < -20.0” --filter-name “ReadPosRankSum-20”.

To maximize the number of SNPs for analyses, we prepared datasets from the genotyped vcf file for each analysis by following filtering using vcftools (Danecek, et al. 2011).

#### Dataset 1: PCA and ADMIXTURE (Fig. 1, Fig. S1)

This dataset includes the individuals listed in Table S2 (dataset1). We removed four shiba dogs, one dingo, and one NGSD from dataset 2. This dataset consisted of 815,014 sites (103 individuals).

#### Dataset 2: *f3*, *f4* statistics (Figs. 3b, 5, S3, S4, S7, S8, S9, S11 and S12)

This dataset includes the individuals listed in Table S2 (dataset 2). We removed sites with missingness higher than 7% and minor allele frequency (MAF) < 0.05. We extracted bi-allelic sites with coverage equal to or more than three in all individuals and with GQ values equal to or more than eight in all individuals. The final dataset consisted of 815,014 sites (109 individuals).

#### Dataset 3: *f3* statistics and *f4*-ratio (Fig. 3a, 5, S5, and S6)

This dataset includes the individuals listed in Table S2 (dataset 3). We removed sites with missingness higher than 30% and minor allele frequency (MAF) < 0.05. We extracted bi-allelic sites with coverage equal to or more than three in all individuals and with GQ values equal to or more than eight in all individuals. The final dataset consisted of 584,646 sites (67 individuals).

#### Dataset 4: Phylogenetic analyses (Figs. S2)

This dataset includes the individuals listed in Table S2 (dataset 4). We removed sites with missing data and minor allele frequency (MAF) < 0.02. We extracted bi-allelic sites with coverage equal to or more than three in all individuals and with GQ values equal to or more than eight in all individuals. After the first filtration, we extracted unlinked biallelic SNPs. The final dataset consisted of 113,784 sites (54 individuals).

#### Dataset 5: Phylogenetic analyses (Fig. 2)

This dataset includes the individuals listed in Table S2 (dataset 5). We removed sites with missing data and minor allele frequency (MAF) < 0.02. We extracted bi-allelic sites with coverage equal to or more than three in all individuals and with GQ values equal to or more than eight in all individuals. After the first filtration, we extracted transversion unlinked biallelic SNPs. The final dataset consisted of 74,761 sites (58 individuals).

#### Dataset 6: *f3* statistics (Fig. S8)

This dataset includes the individuals listed in Table S2 (dataset 6). We removed sites with missingness higher than 20% and minor allele frequency (MAF) < 0.05. We extracted bi-allelic sites with coverage equal to or more than three in all individuals and with GQ values equal to or more than eight in all individuals. After the first filtration, we extracted transversion SNPs. The final dataset consisted of 725,285 sites (54 individuals).

### Phylogenetic analysis

We extracted unlinked biallelic SNPs using PLINK ver. 1.9 (Purcell, et al. 2007) with an option “--indep-pairwise 50 10 0.1”. The pruned SNP vcf file was converted to PHYLIP format. 10 kb sequences from the 5’ end of the PHYLIP format file were extracted and a model for the Maximum Likelihood method was selected using MEGA ver. X (Kumar, et al. 2018). A phylogenetic tree was constructed using the Maximum Likelihood (ML) method using PhyML ver. 3.2 (Guindon, et al. 2010) with a model selection option “-m GTR” and with 100 bootstrap replications.

### Principal component analysis and ADMIXTURE

We performed a principal component analysis (PCA) using PLINK ver. 1.9 (Purcell, et al. 2007) with an option “--indep-pairwise 50 10 0.1” to explore the affinity among canids (Figure 1a).

ADMIXTURE ver. 1.3 (Alexander and Lange 2011) was run on the dataset 1 (Fig. 1b and Fig. S1) assuming 2 to 8 clusters (K=2-8).

### *f3*, *f4* statistics, and *f4*-ratio

*f*3, *f4* statistics, and *f4*-ratio implemented in ADMIXTOOLS ver. 7.0.1 (Patterson, et al. 2012) were used to evaluate the shared genetic drift among gray wolves, Japanese wolves, and modern and ancient dogs using SNP dataset 2, 3, and 6.

### qpWave and qpGraph

We used qpWave as implemented in Admixtools2 (Maier, et al. 2022) to study the number of admixture flows from wolves to dogs. We set the test populations (dogs) and the source populations as describe in Table 1.

We used qpGraph as implemented in Admixtools2 (Maier, et al. 2022) to compute the admixture graph with best fitting model by adding one group after the other and compared the likelihood scores. We started from (outgroup: coyote(Eurasian wolf: wolf chanco(West Eurasian dog: African dong, East Eurasian dog: NGSD))), and added Japanese wolf (Jw284), Jomon dog (MD1), and Greenland sled dog, one by one.

### Availability of data

The nucleotide sequences were deposited in the DDBJ nucleotide database under Bioproject PRJDB13874.

## Acknowledgments

We would like to express our gratitude to all the participants who contributed to this study. Sequencing of the samples in this study was supported by JSPS KAKENHI Grant Number 16H06279 (PAGS). This study was supported by Grant-in-Aid for Scientific Research (22H00737, 22H00013, 20K01104, 19H0053), Collaborative research grant, Evolutionary Studies of Biosystems (ESB), The Graduate University for Advanced Studies (Sokendai) (2020, 2021) and Student Dispatch Program, Sokendai (2022).

## Author contribution

Yohey Terai *(concept, , analyses, write ms, construction of libraries, sequencing)

Xiayire Xiaokaiti, (part of analysis, DNA extraction, construction of libraries)

Jun Gojobori, (part of analysis, edit ms)

Nami Arakawa, (construction of libraries)

Takao Sato, (provided samples, archaeological information)

Kenji Kasai, (provided samples, archaeological information)

Kenichi Machida, (provided samples, archaeological information)

Kyomi Yamazaki, (provided samples, archaeological information)

Naomitsu Yamaji, (provided samples, archaeological information)

Hitomi, Hongo, (archaeological information, edit ms)

Takashi Gakuhari, (edit ms, DNA extraction, construction of libraries)

