## Supplemental Figures for "Genome analysis of the Jomon dogs reveals the oldest domestic dog lineage in Eastern Eurasia"

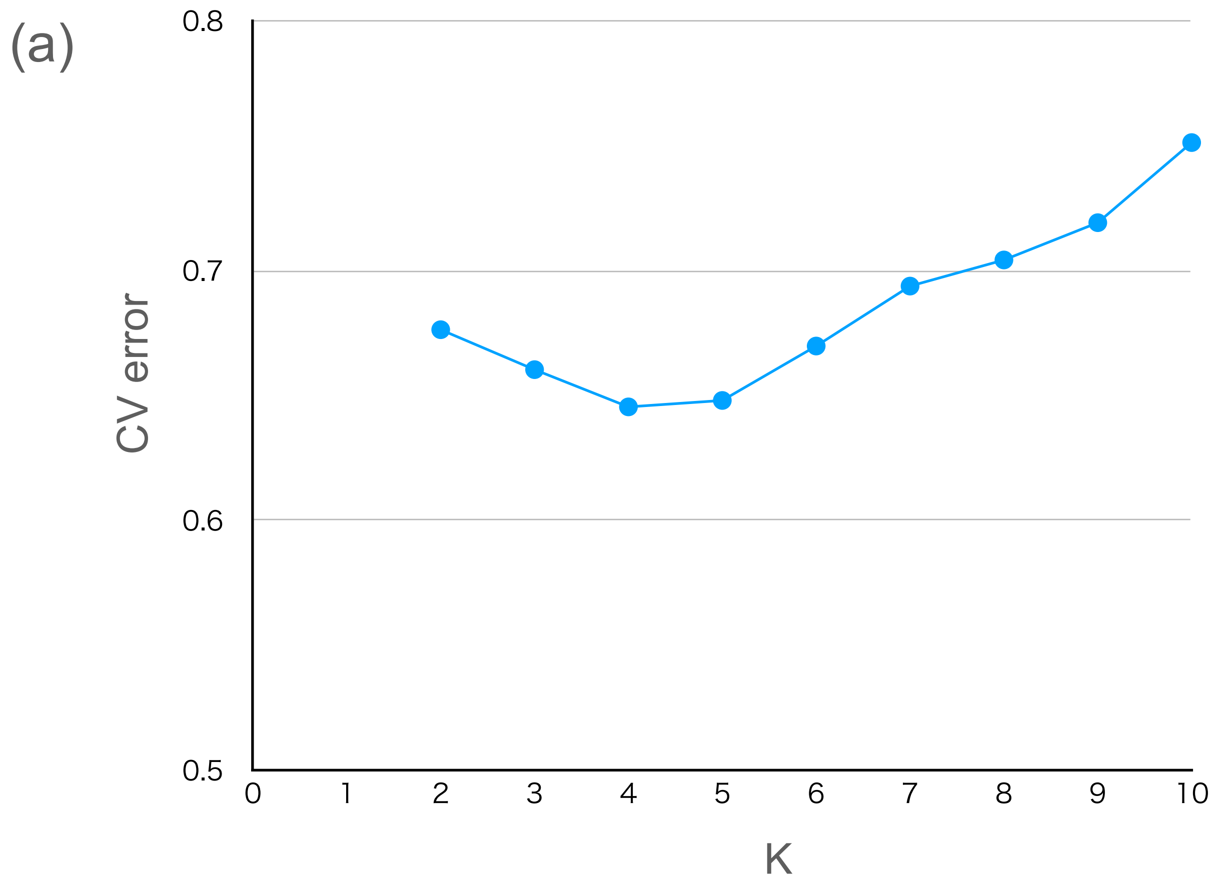


Figure S1
(a) Cross validation (CV) values for ADMIXTURE analysis of SNP data. (b) ADMIXTURE results based on SNP data for K = 2-8 (see Table S1 for sample information).

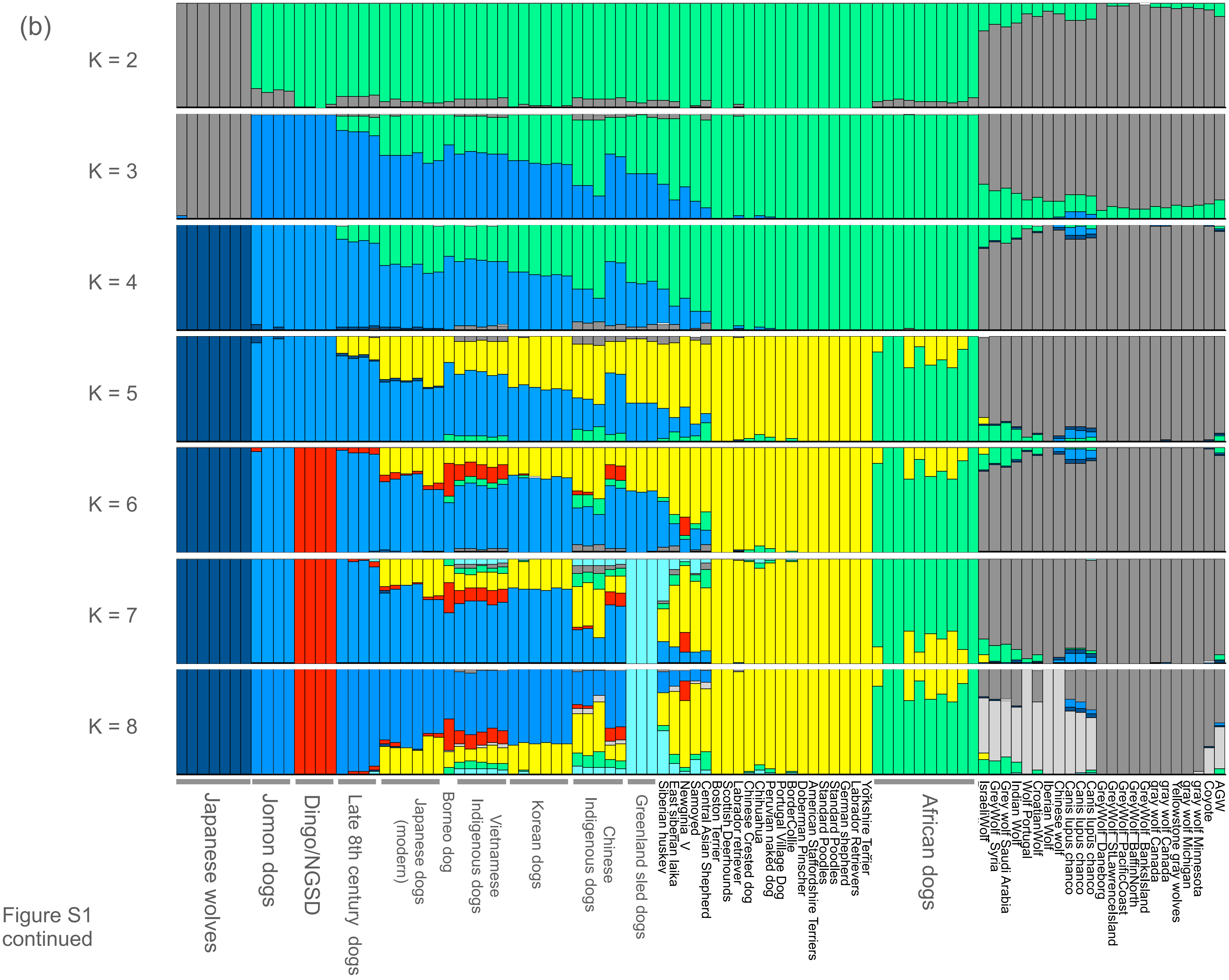


Figure S1 continued

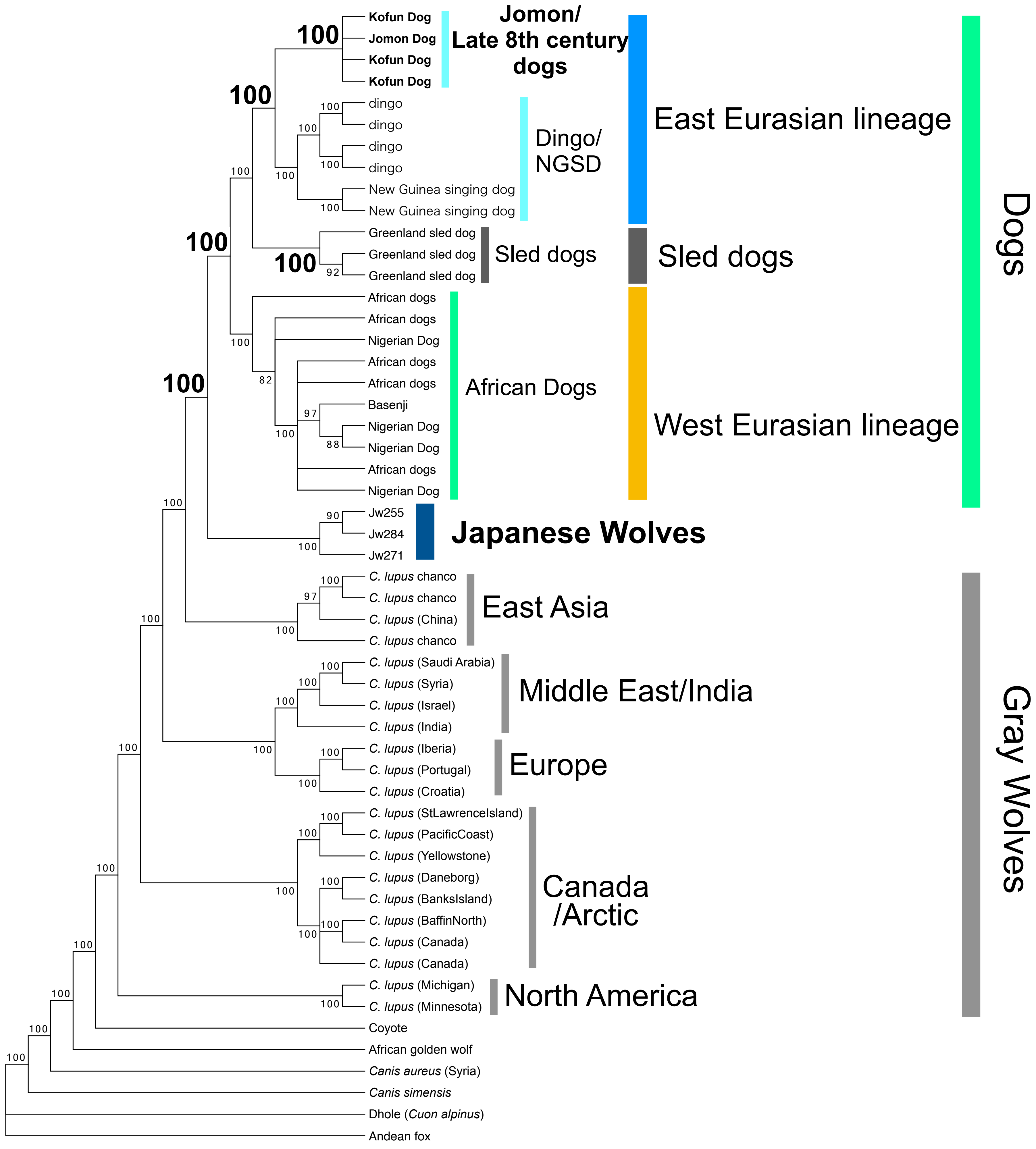


Figure S2
Maximum likelihood tree based on 113,784 unlinked biallelic SNPs. Node labels indicate bootstrap replicates.

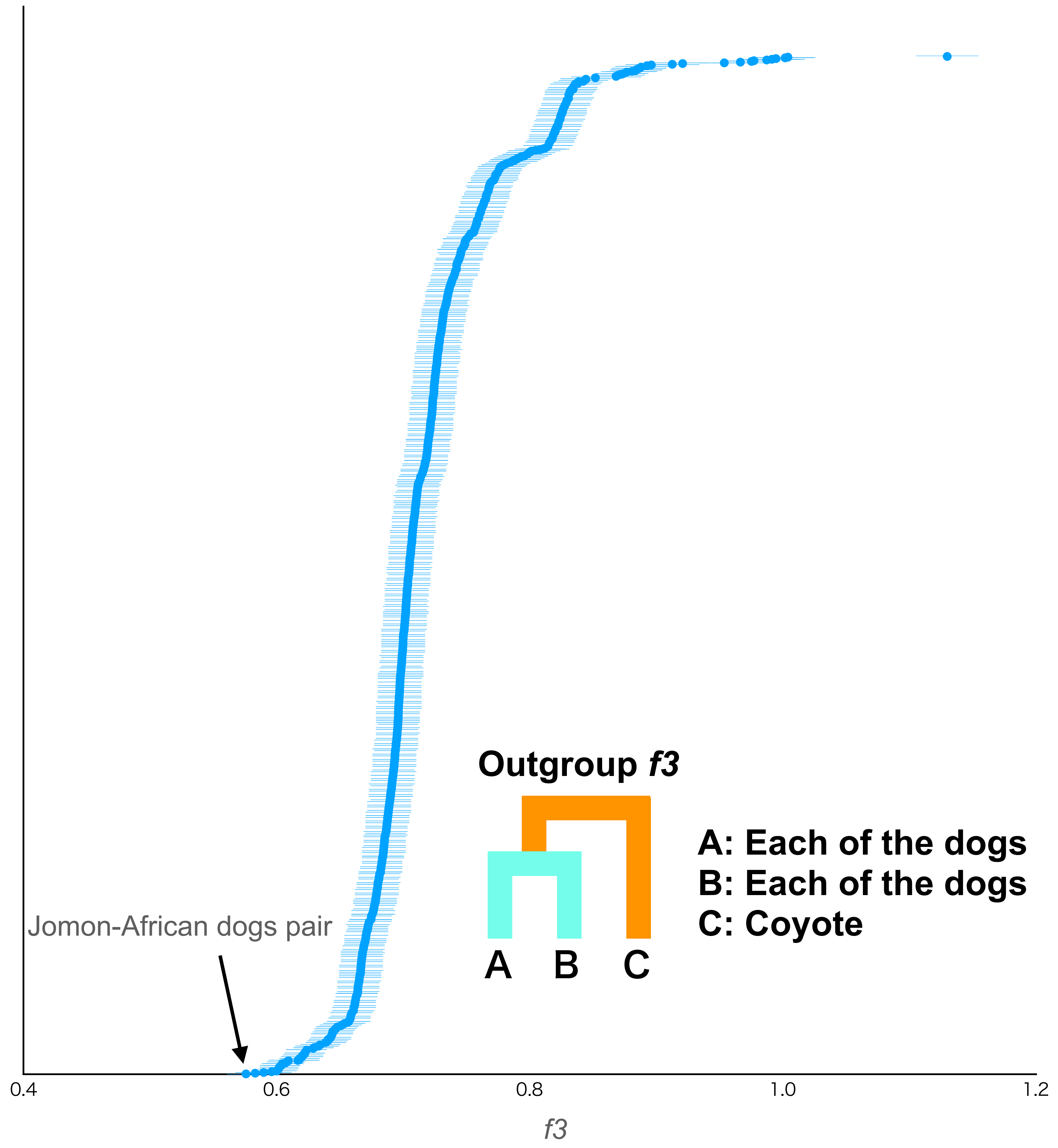


Figure S3
Shared genetic drift between all dog combinations measured by outgroup *f3* statistics. African dog individuals were used as populations. Each *f3* statistical value is plotted in order of highest to lowest value from the top. Error bars represent standard errors. In 1431 combinations, the pair with the smallest *f3* value was a Jomon (MD1) and African dogs pair.

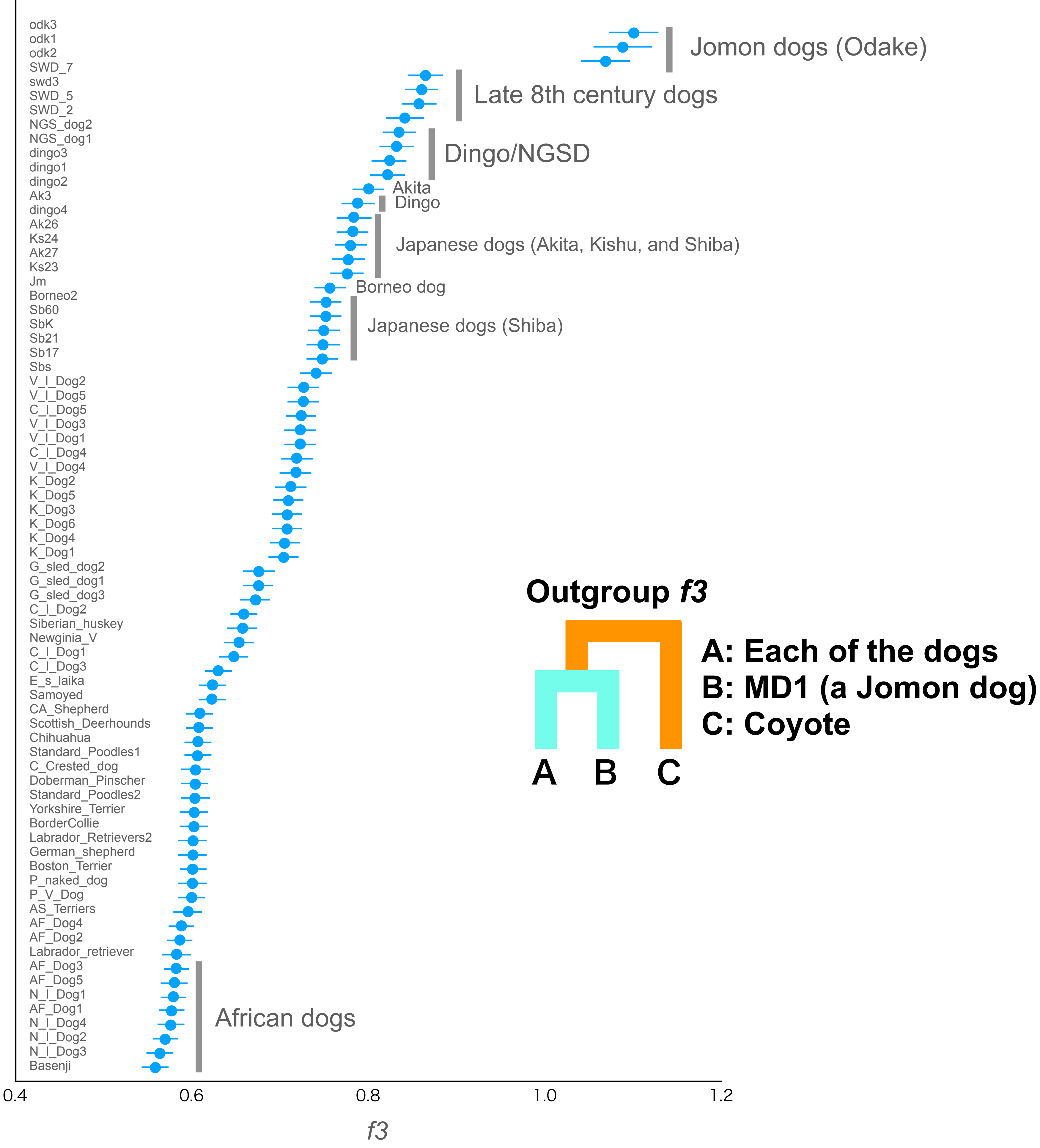


Figure S4
Shared genetic drift between a Jomon dog (MD1) and all other dogs measured by outgroup *f3* statistics. Each *f3* statistical value is plotted in order of highest to lowest value from the top, and the names of the wolves are shown on the left side of each panel. Error bars represent standard errors.

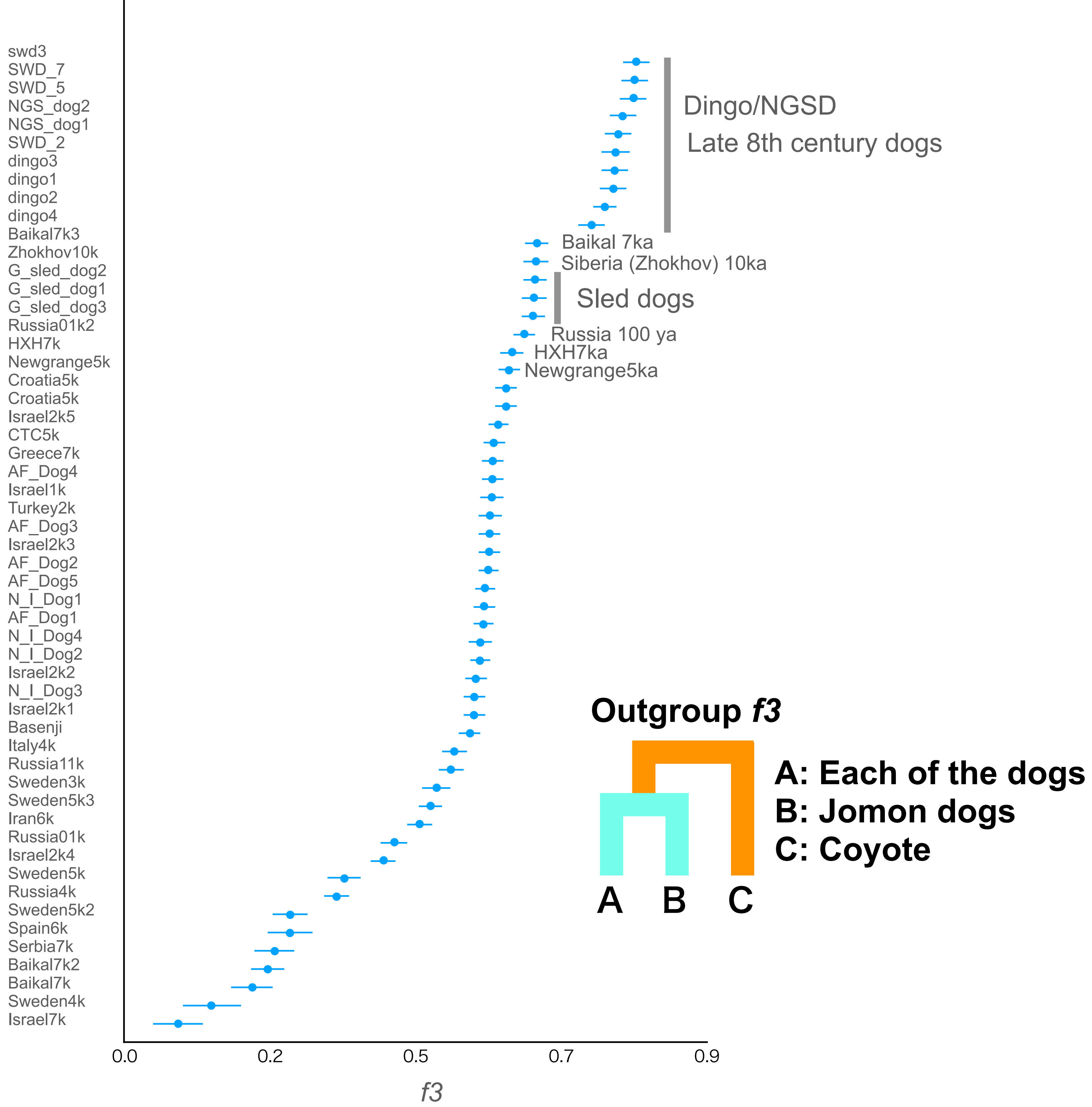


Figure S5
Shared genetic drift between a Jomon dog (MD1) and all other dogs measured by outgroup *f3* statistics. The Jomon dog individuals were used as a population. Each *f3* statistical value is plotted in order of highest to lowest value from the top, and the names of the wolves are shown on the left side of each panel. Error bars represent standard errors.

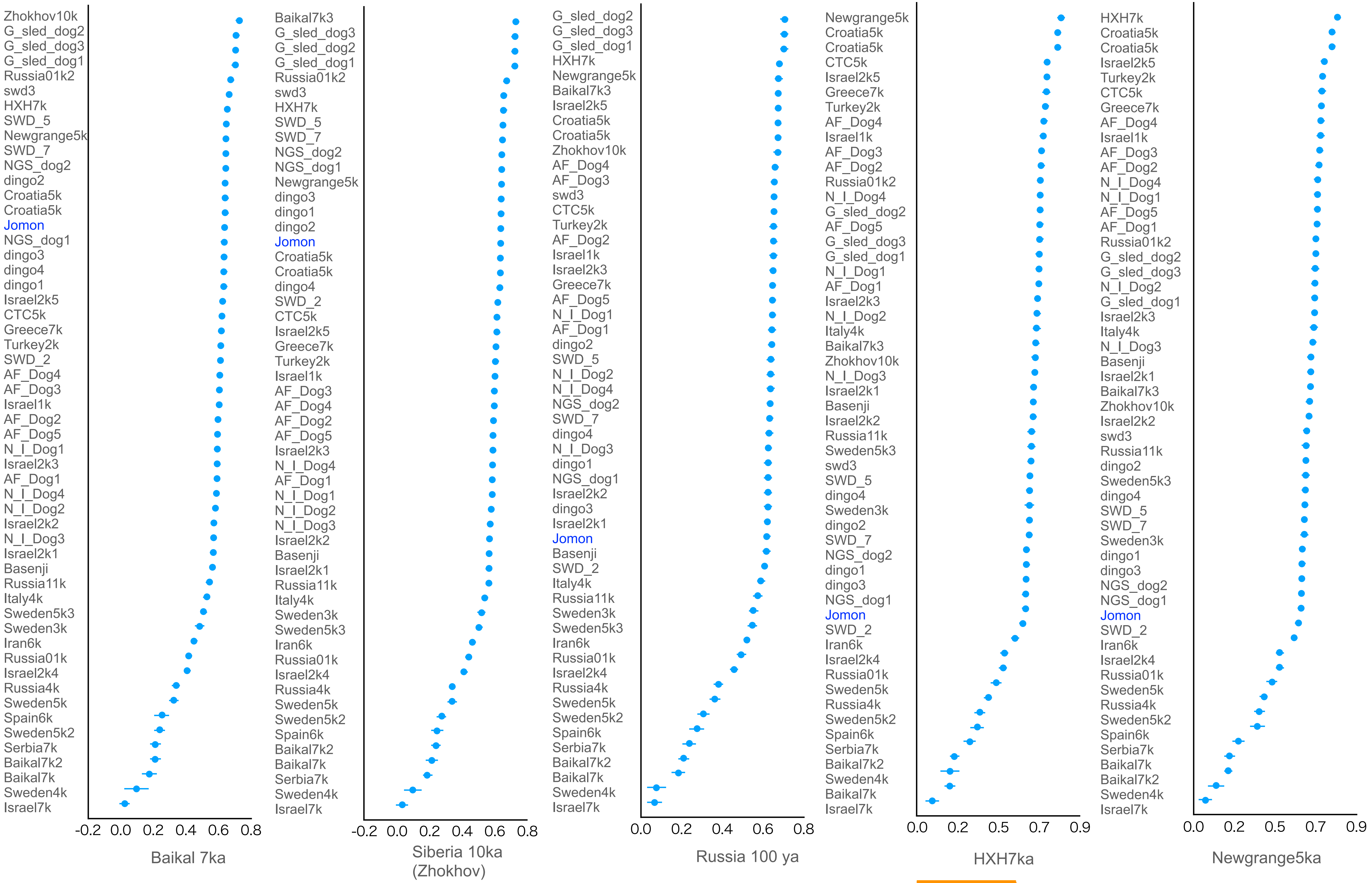


Figure S6
Shared genetic drift between ancient dogs (under each panel) and all other dogs measured by outgroup *f3* statistics. The Jomon dog individuals were used as a population. Each *f3* statistical value is plotted in order of highest to lowest value from the top, and the names of the wolves are shown on the left side of each panel. Error bars represent standard errors.

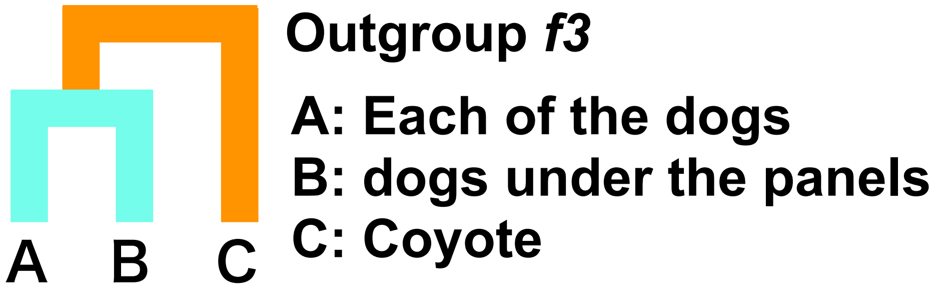

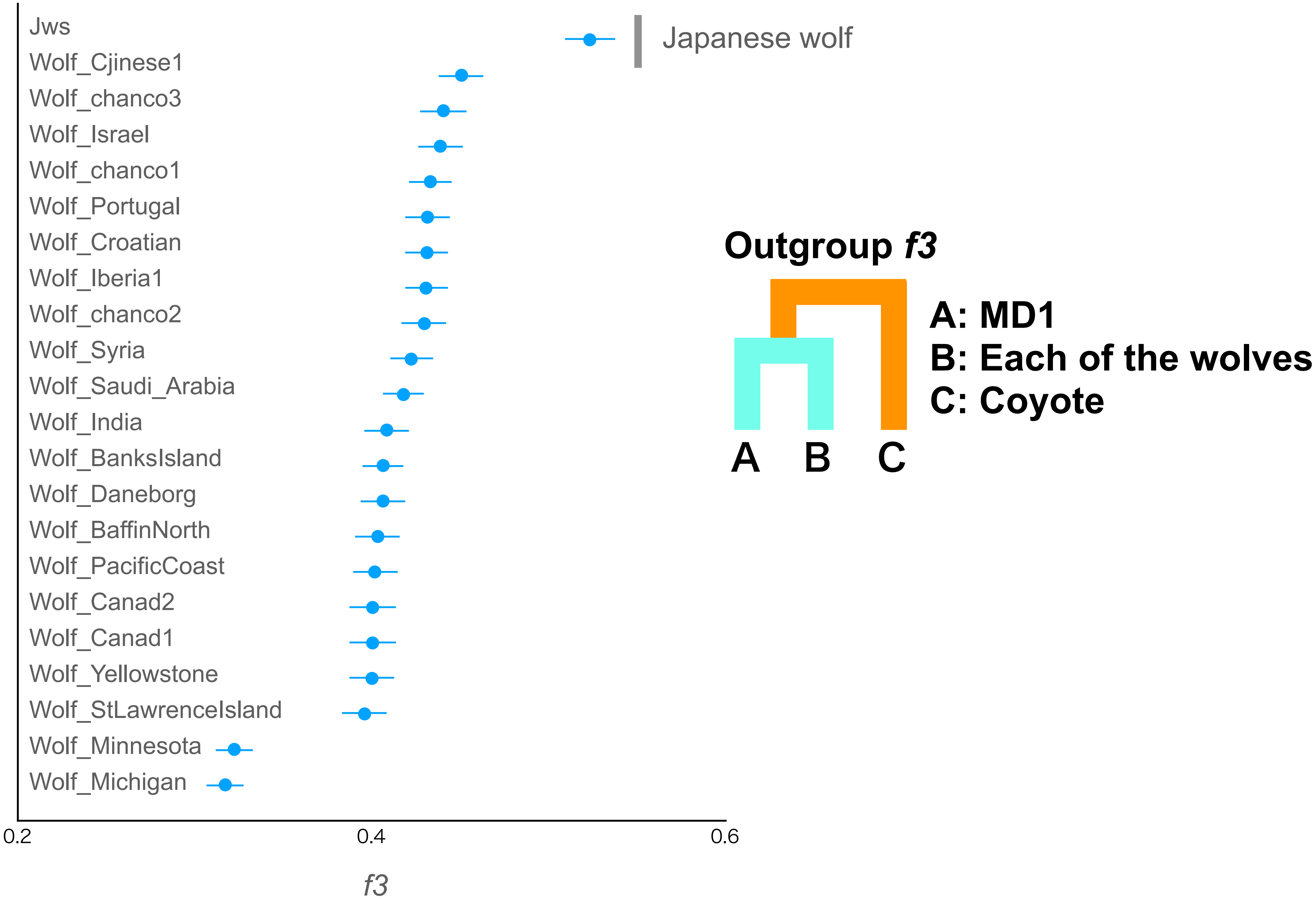


Figure S7
Shared genetic drift between a Jomon dog (MD1) and all other wolves measured by outgroup *f3* statistics. Each *f3* statistical value is plotted in order of highest to lowest value from the top, and the names of the wolves are shown on the left side of each panel. Error bars represent standard errors.

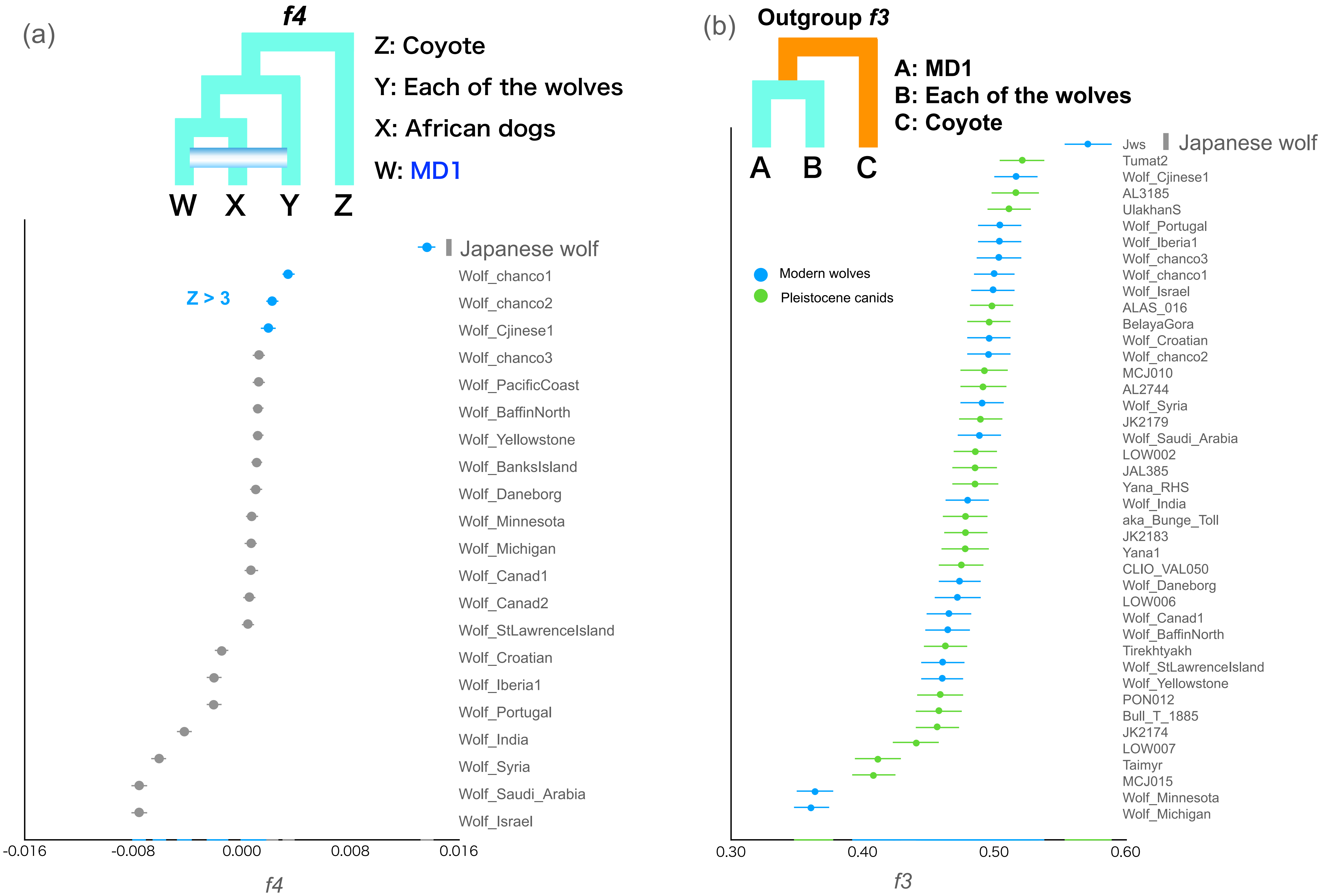


Figure S8

(a) *f4* statistics testing the genetic affinity of a Jomon dog (MD1) with all other wolves. *f4* values for each combination are plotted. We computed *f4* statistics where Y in the schematic representation represents the other dogs. Each *f4* value is listed in order of highest to lowest value from the top.(b) Shared genetic drift between a Jomon dog (MD1) and all other wolves measured by outgroup *f3* statistics. Each *f3* statistical value is plotted in order of highest to lowest value from the top. Error bars represent standard errors. The names of the wolves are shown on the right sides of the panels.

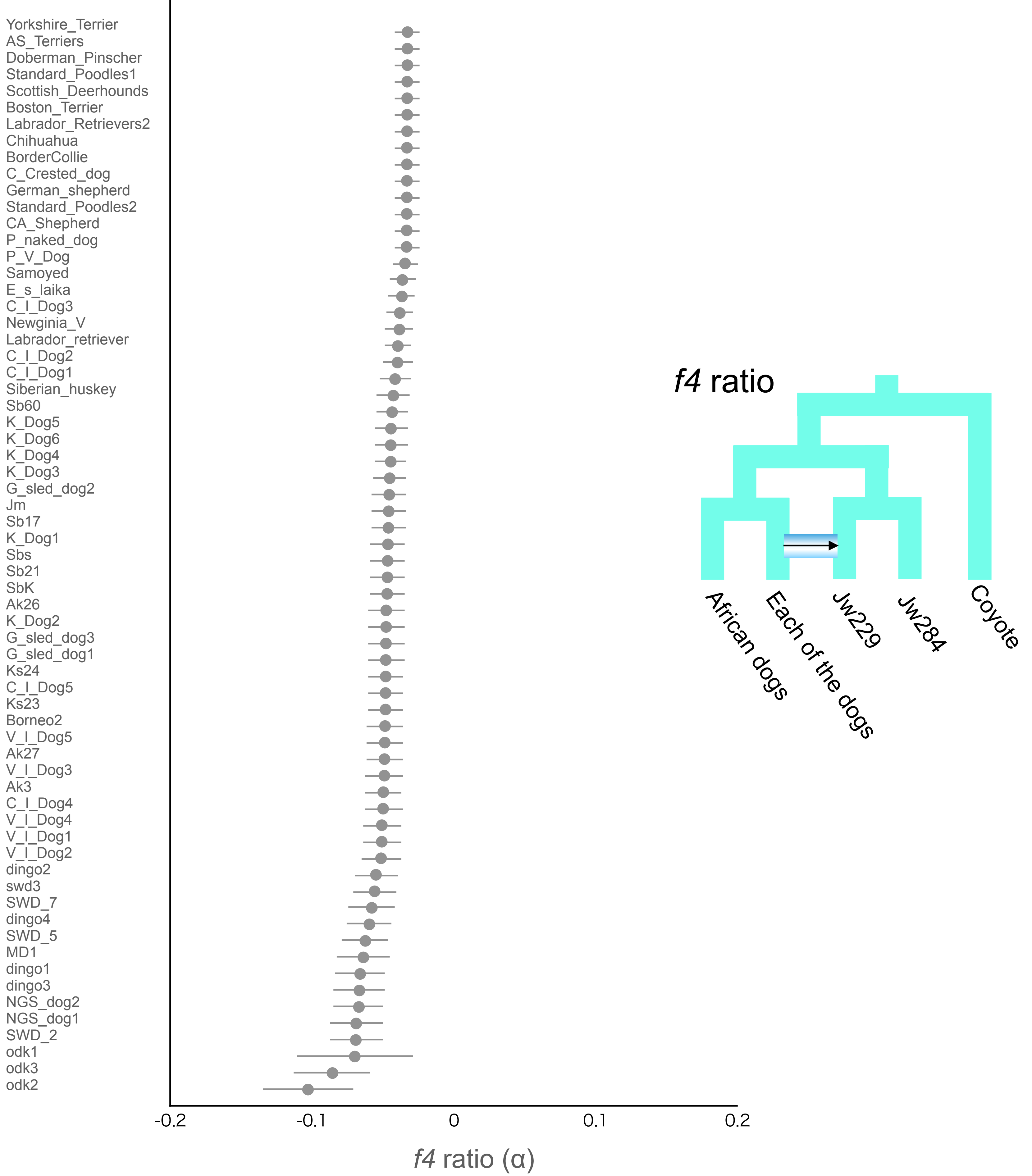


Figure S9

*f4*-ratio test to estimate proportion of genome introgression from dogs to the Japanese wolf. Each *f4*-ratio α value is plotted in order of lowest to highest value from the top, and the names of the dogs are shown on the left side of the panel. Error bars represent standard errors.

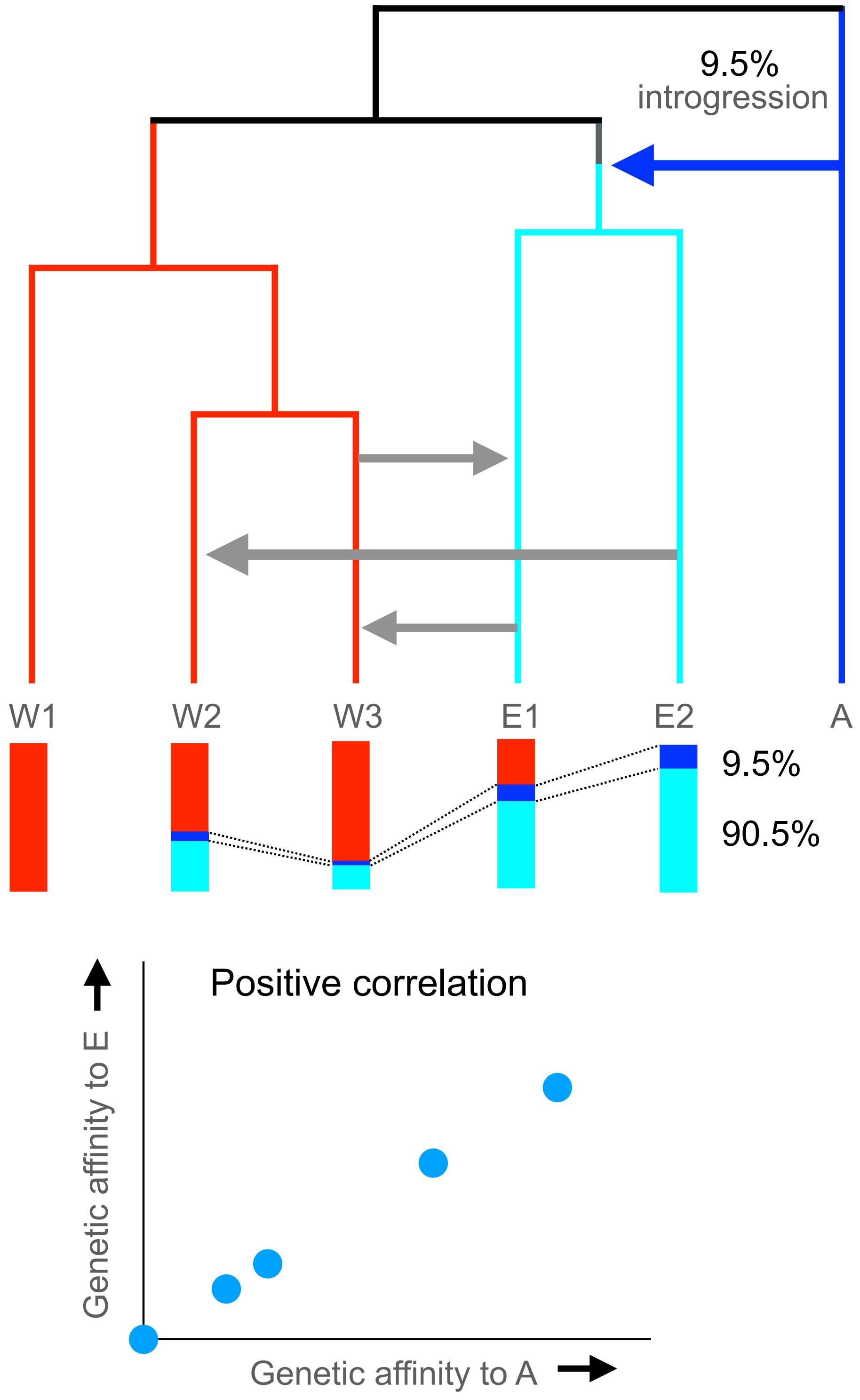


Figure S10. Models of a single introgression event and diffusion of the introgressed genome.
Upper left panel: if there was a single event of introgression from ancestral lineage A (shown in blue) to lineage E (shown in light blue), for example, 9.5% of the genome introgressed from lineage A to lineage E (A). After the first introgression, the genome of lineage E contains 9.5% of the genome of lineage A. When the genome of lineage E introgresses to lineage W, the ratio of the genomes of E to A (E: 90.5%, A: 9.5%) is maintained in the genome of lineage W.
Conversely, when the genome of lineage W introgresses to lineage E, the ratio of the genomes of lineage E and A is maintained. Therefore genetic affinity with A positively correlates with genetic affinity with E in all individuals, lower panel.

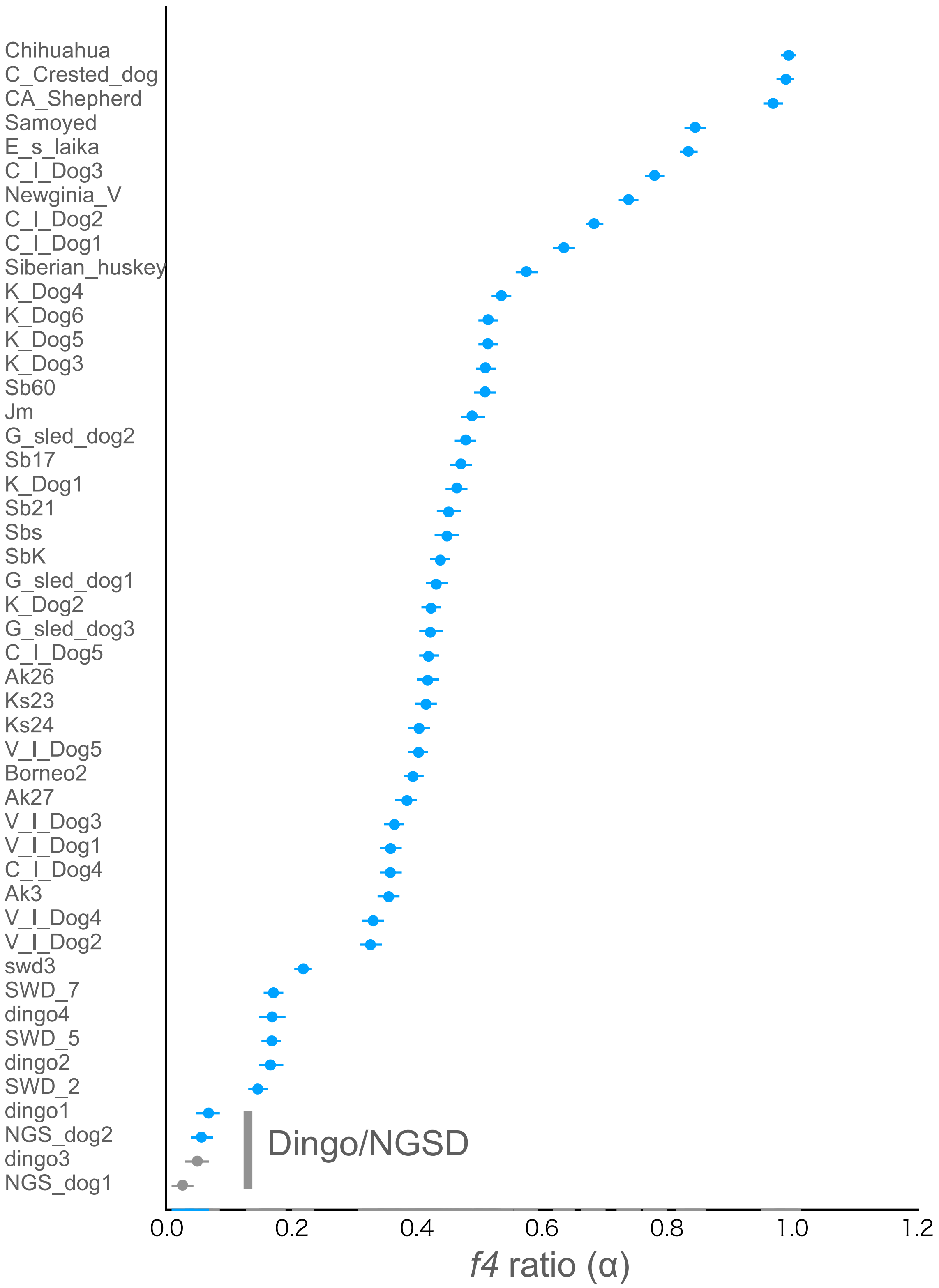

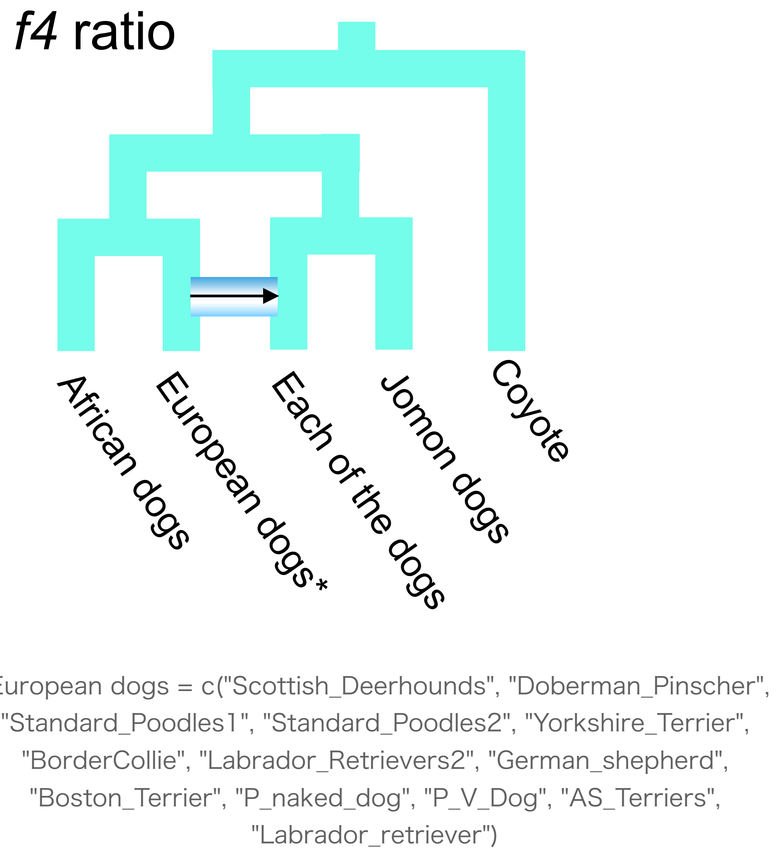


Figure S11

*f4*-ratio test to estimate proportion of genome introgression from the European dogs to East Eurasian dogs. Each *f4*-ratio α value is plotted in order of highest to lowest value from the top, and the names of the dogs are shown on the left side of the panel. Error bars represent standard errors. Z score above 3 is colored in blue.

dataset2

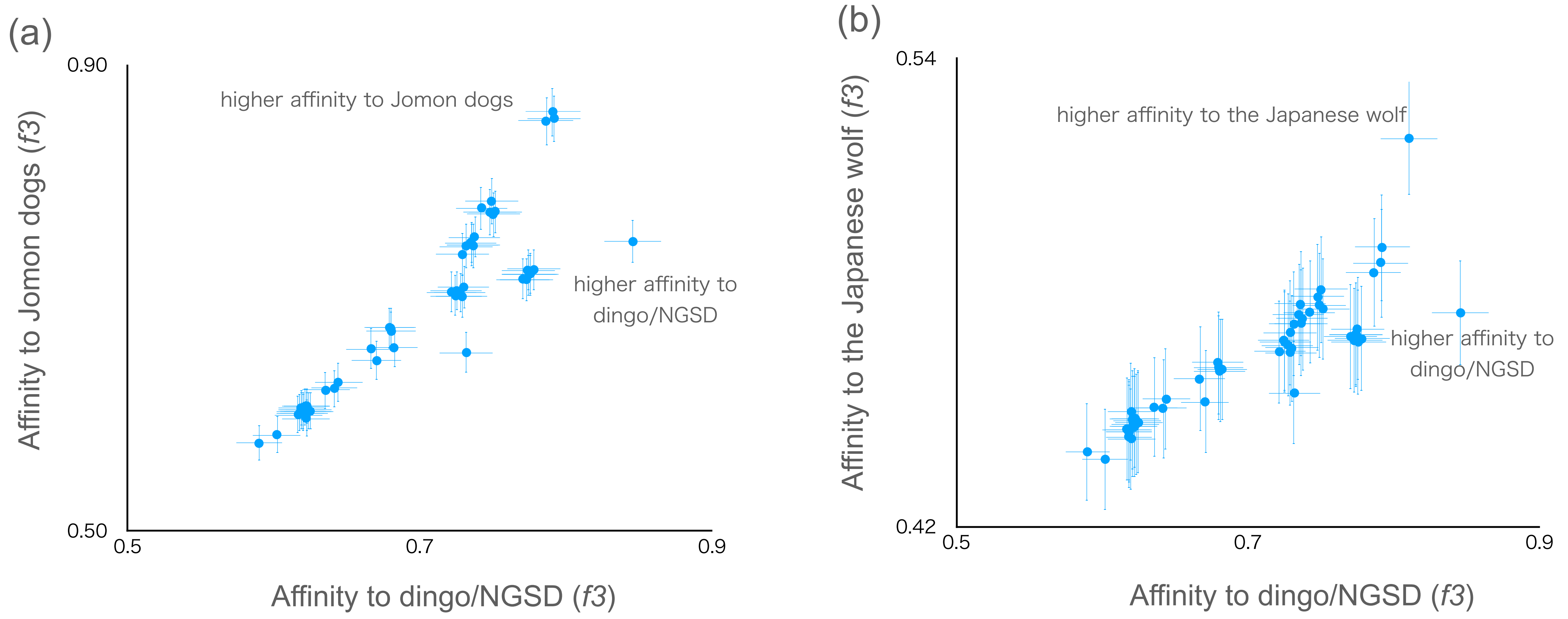


Figure S12

*f3* statistics testing whether dogs share more alleles with dingo/NGSD (x-axis) or Jomon dogs (a) and Japanese wolf (b) (y-axis). Dots show the *f3* statistics, and horizontal and vertical error bars represent standard errors.

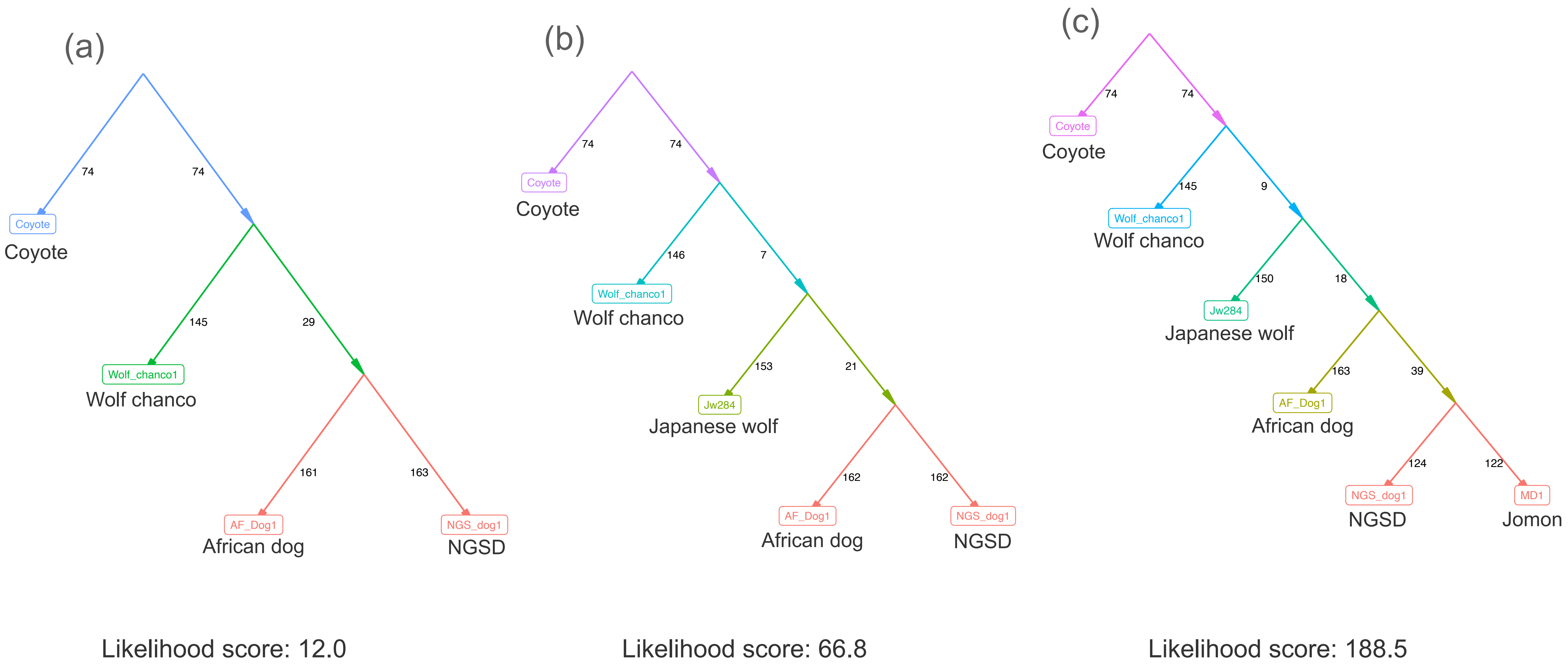


Figure S13 Admixture graph modelings
We started building graphs consisting of coyote, wolf chanco, African dog, and NGSD (a) and added the remaining samples sequentially in the following order: (b) Japanese wolf, (c) Jomon dog. We added an introgression from the ancestor of Japanese wolf to ancestor of Jomon dogs (d) and further an admixture event between Jomon and No-Jw-dogs (e). Likelihood scores are shown under the models.

(d)

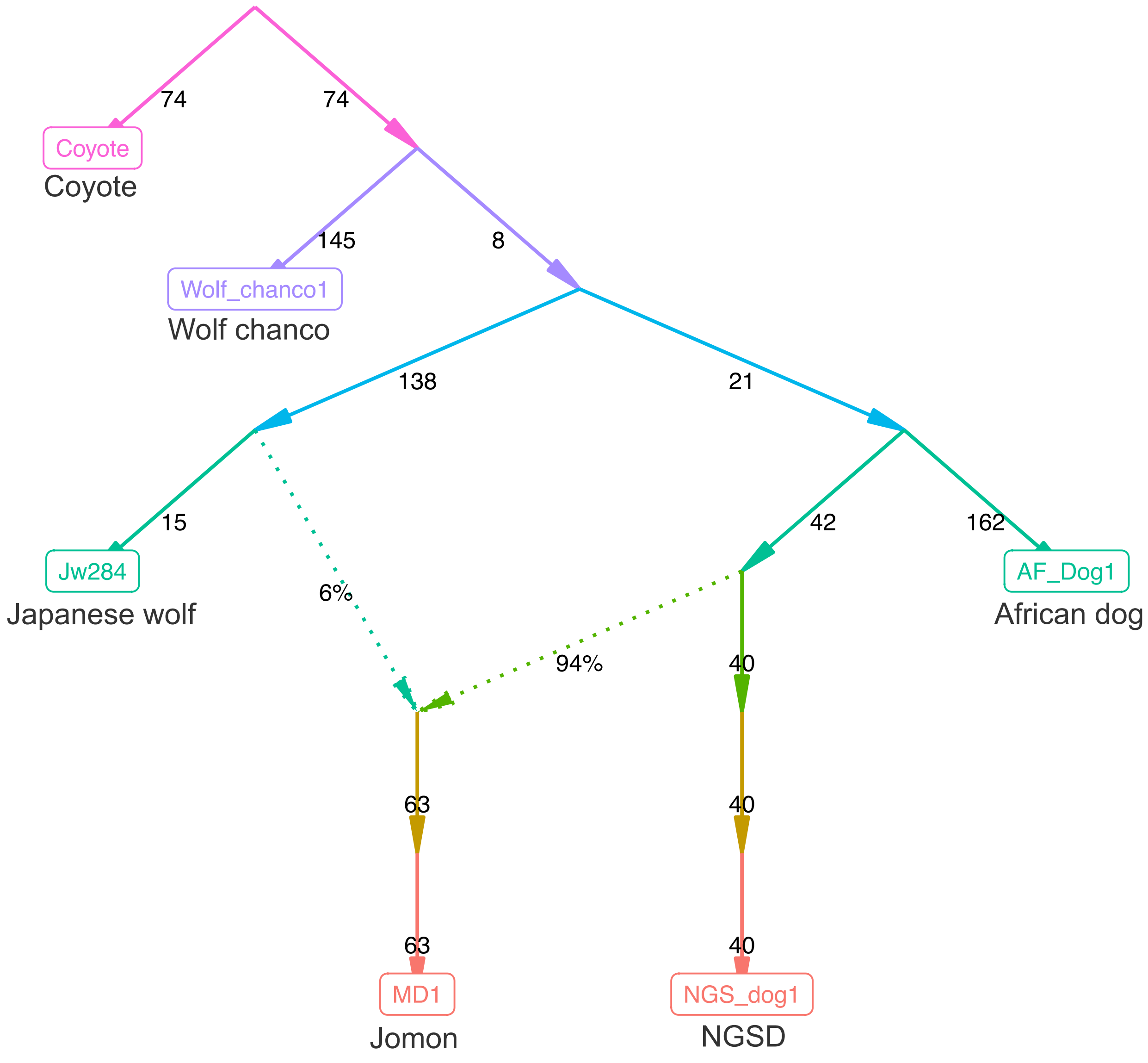


Likelihood score: 80.2

(e)

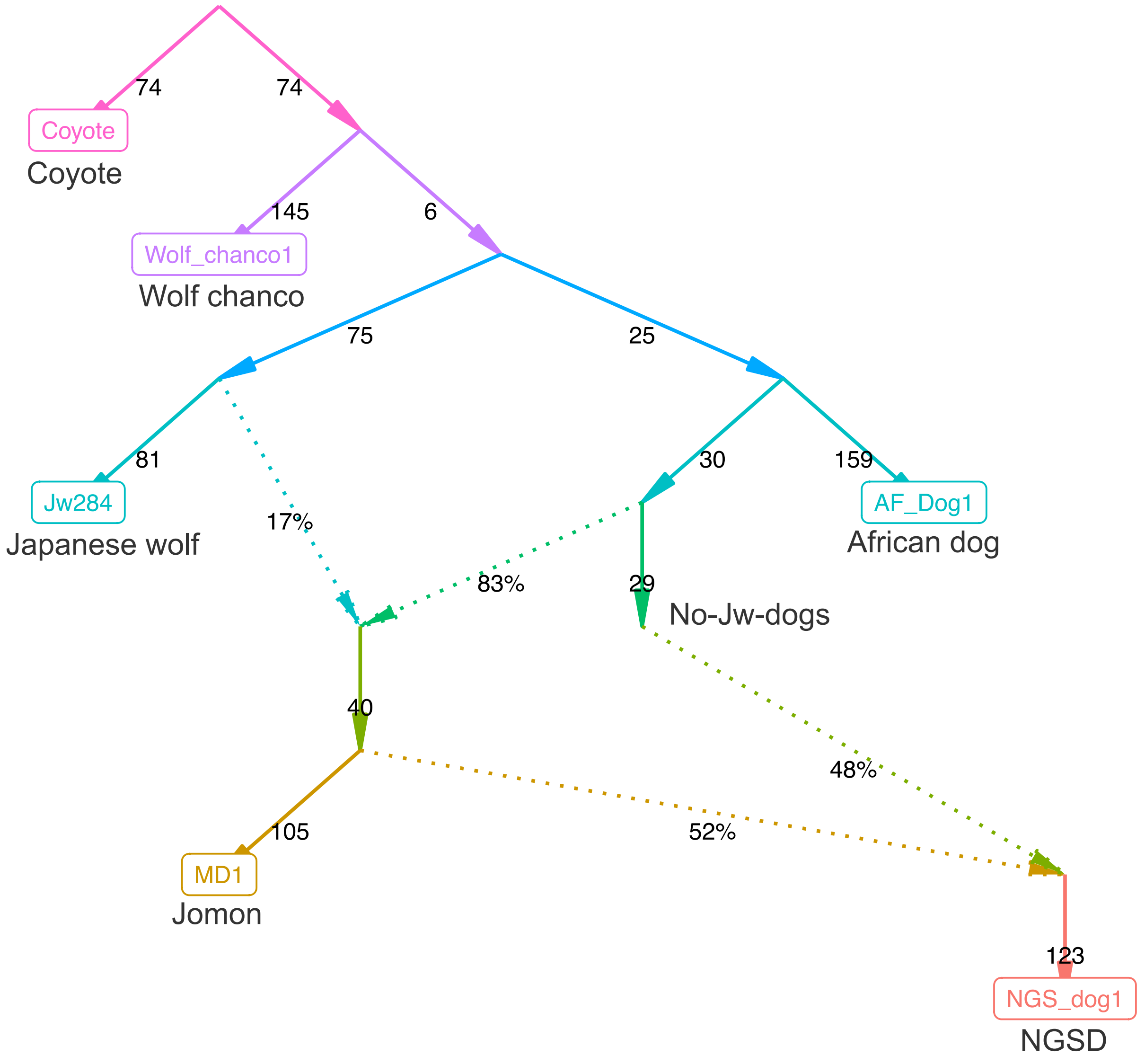


Likelihood score: 29.0

Figure S13 continued

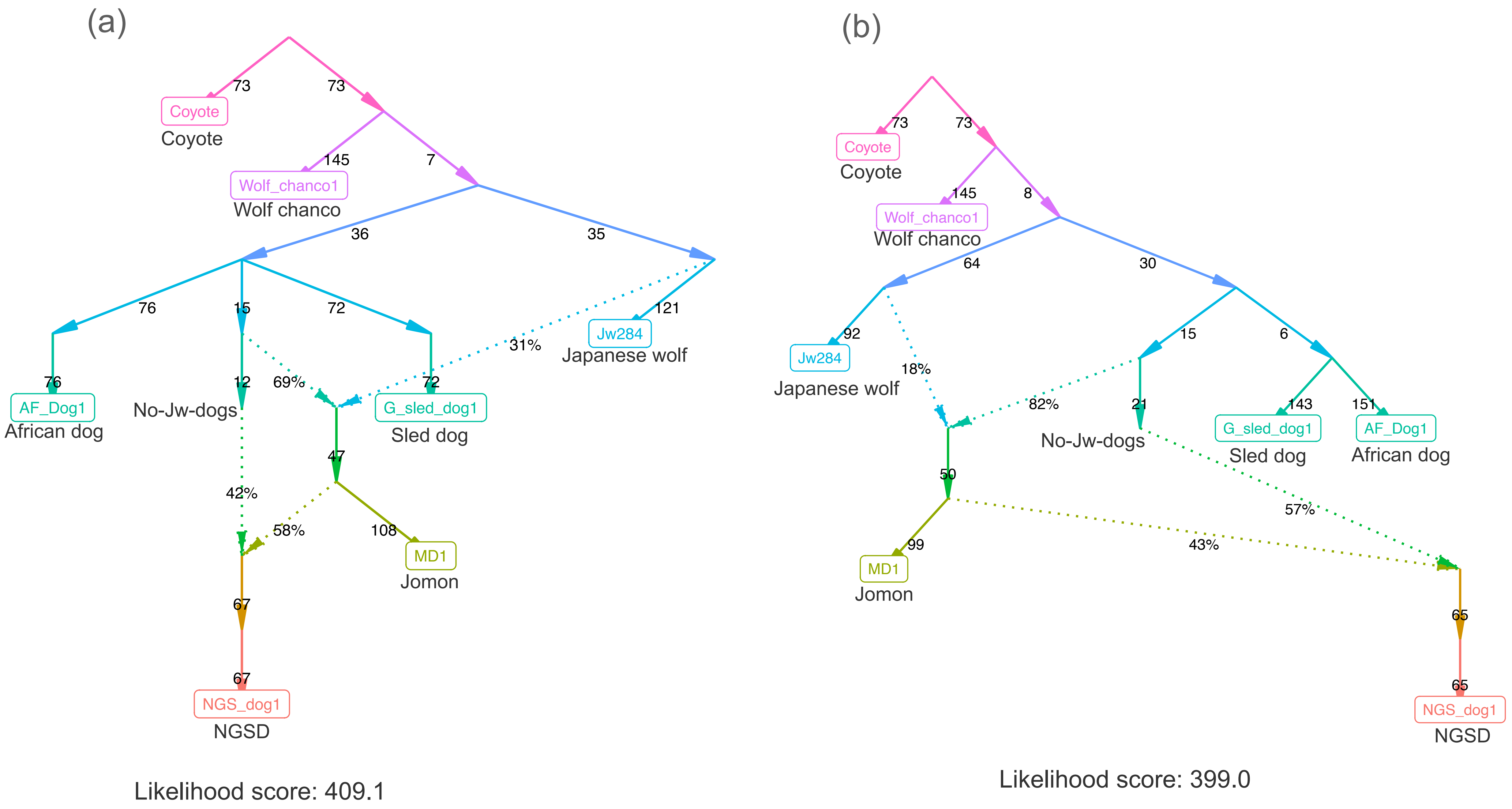


Figure S14 Admixture graph modelings

In addition to a model in Fig. S13e (a), we added a Greenland sled dog. The models represent that sled dog lineage has diverged (a) simultaneously with the divergence of Eastern and Western Eurasian lineages, (b) from the Western Eurasian lineage, (c) from the dingo/NGSD lineage, (d) from the Jomon dog lineage, or have arisen by the admixture of (e) the Jomon dog and Western Eurasian lineages, (f) the No-Jw-dog and Western Eurasian lineages, and (g) the dingo/NGSD and Western Eurasian lineages. Likelihood scores are shown under the models.

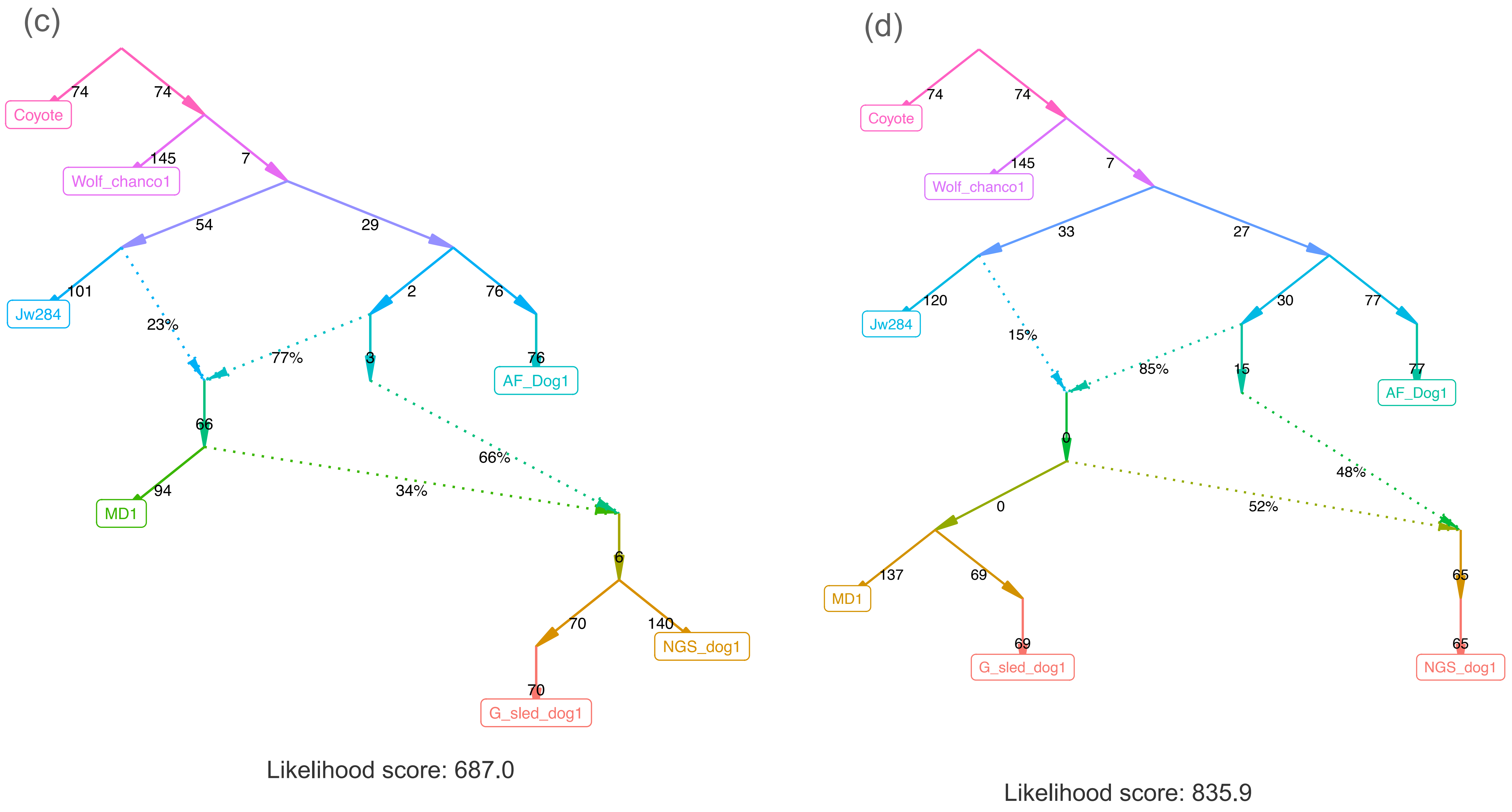


Figure S14 continued

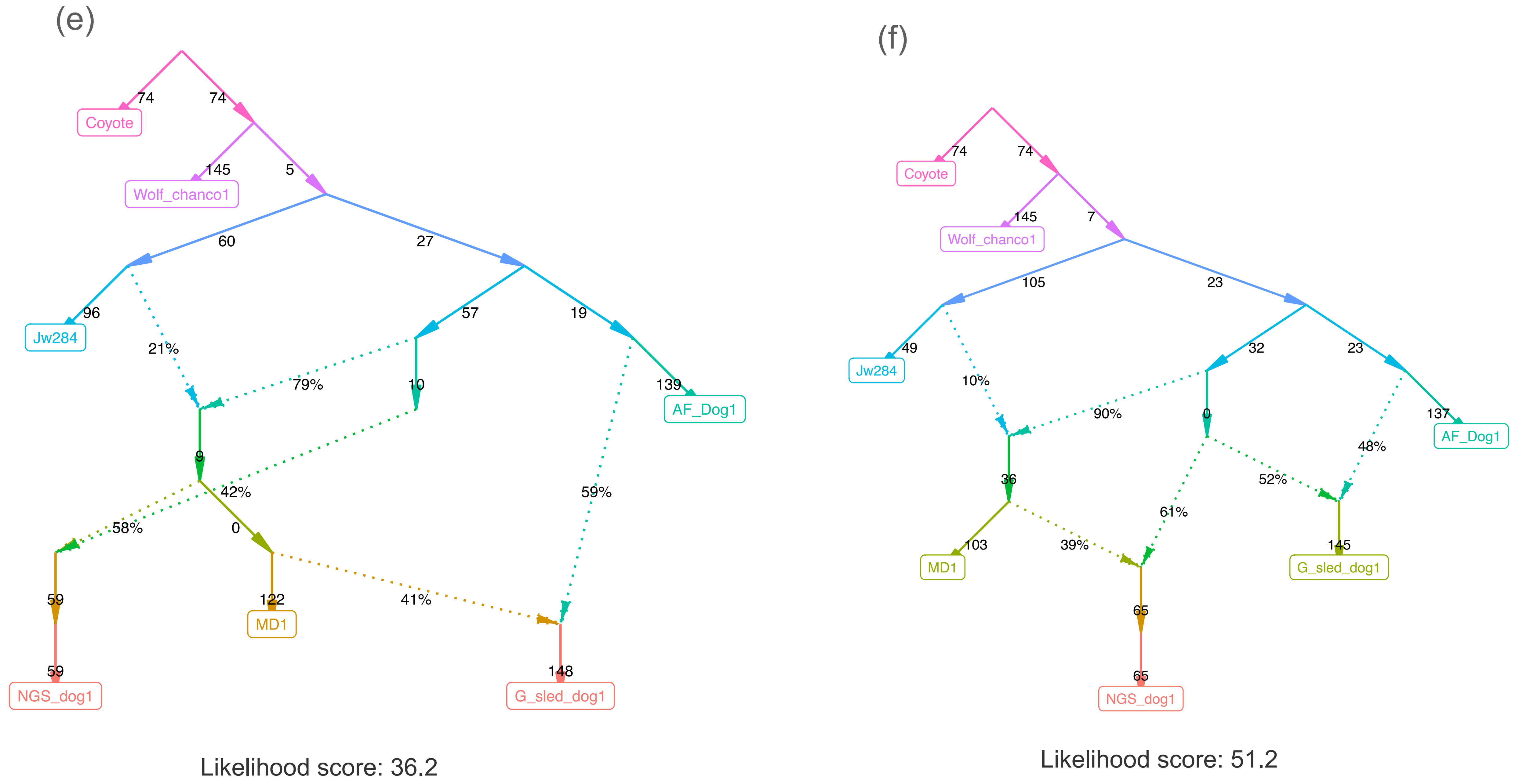


Figure S14 continued

(g)

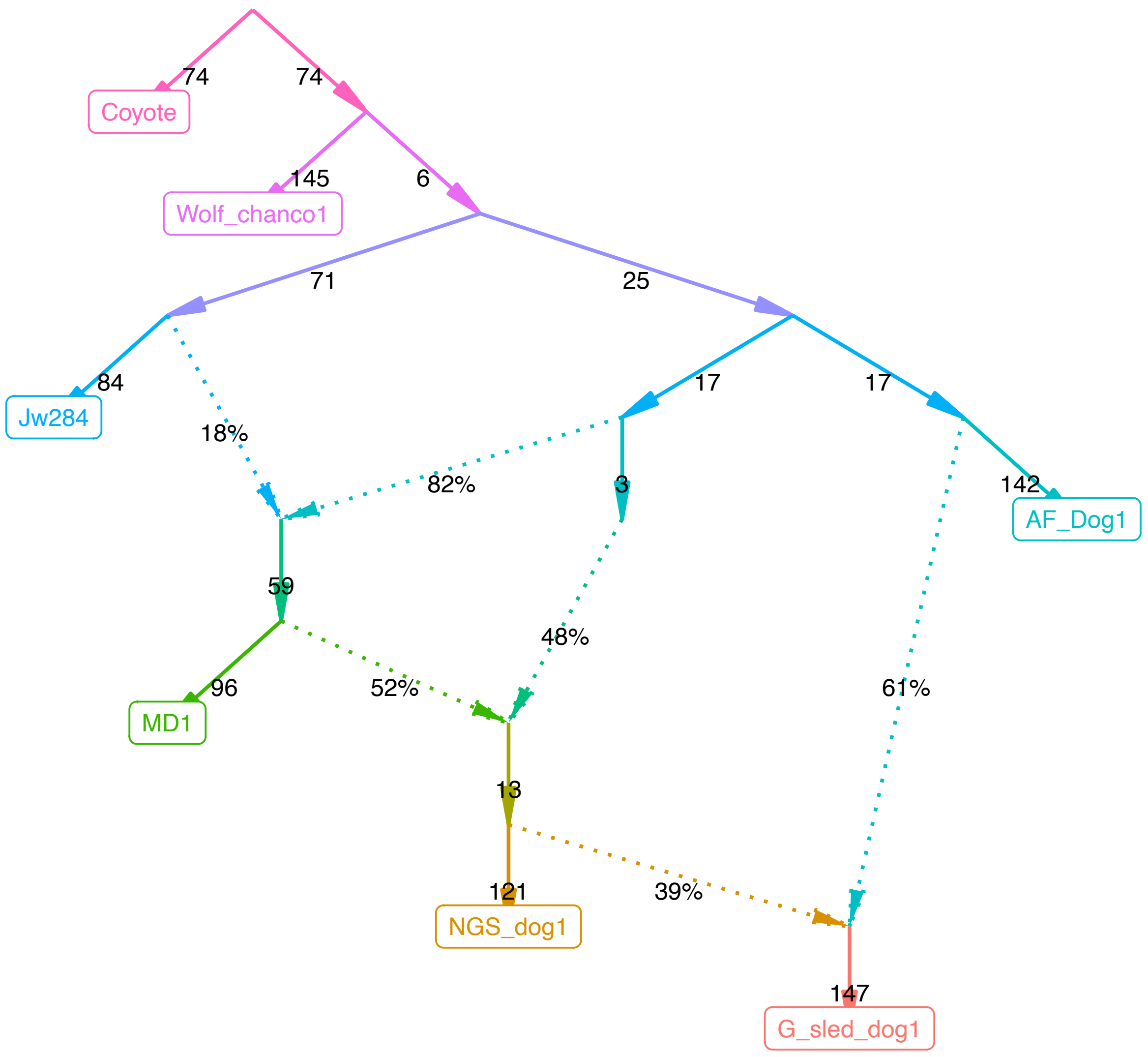


Likelihood score: 31.7

Figure S14 continued
