## Supplemental Table S1 for "Genome analysis of the Jomon dogs reveals the oldest domestic dog lineage in Eastern Eurasia"

### Table S1 Determined sequences in this study

| ID | Period | Isolation site (Prefecture) | Total mapped reads | Average coverage | Reference covered |
|---|---|---|---|---|---|
| odk1 | Jomon | Toyama Pref. | 21,.0 Gb | 8.6x | 11% |
| odk2 | Jomon | Toyama Pref. | 15.1 Gb | 6.2x | 45% |
| odk3 | Jomon | Toyama Pref. | 44.1 Gb | 18.2x | 19% |
| MD1 | Jomon | Chiba Pref. | 18.3 Gb | 7.5x | 80% |
| SWD2 | late 8th century | Chiba Pref. | 15.5 Gb | 6.4x | 75% |
| SWD3 | late 8th century | Chiba Pref. | 96.8 Gb | 40.0x | 96% |
| SWD5 | late 8th century | Chiba Pref. | 23.4 Gb | 9.7x | 89% |
| SWD7 | late 8th century | Chiba Pref. | 23.9 Gb | 9.9x | 86% |
